# Small Extracellular Vesicles from Gram-Negative Bacterial Infections Induce Myeloid Differentiation via p38 Signaling and Coordinate Antibacterial Defense Through the p38–IL-6 Axis

**DOI:** 10.64898/2025.12.30.697125

**Authors:** Adam Fleming, Heather Hobbs, Graham Matulis, Weidong Zhou, Valerie Calvert, Hunter Mason, Samson Omole, Shivam A. Gaikwad, Cameron C. Ellis, Emanuel F. Petricoin, Igor C. Almeida, Ramin M. Hakami

## Abstract

Much still remains to understand about the underlying molecular mechanisms by which the trafficking of small extracellular vesicles (sEVs) modulates innate immune responses during infection with pathogenic Gram-negative bacteria. To address this significant gap in knowledge, we used two infection models to investigate innate immune regulation by the sEVs released from cells infected with either *Yersinia pestis* (Yp) or *Burkholderia thailandensis* (Bt), designated as EXi-Yp and EXi-Bt respectively. The EXi induced differentiation of naïve human monocytes to macrophages and triggered robust pro-inflammatory cytokine release, including release of IL-6, mirroring direct bacterial infection effects. Comprehensive cell signaling analyses revealed that the EXi modulate a small set of host signaling proteins, with p38 activation being primarily responsible for the observed protective effects. EXi-induced p38 activation leads to increased IL-6 release, which in turn is responsible for decreased bacterial survival within recipient immune cells that are subsequently infected. Consistent with the *in vitro* results, mice administered with EXi-Yp exhibited elevated serum IL-6 levels and were protected from Yp infection. Furthermore, using our microfluidic chip platform that allows functional interrogation of EV effects under physiologically relevant conditions, we have demonstrated that EXi exchange between Yp-infected cells and naive recipient monocytes leads to differentiation of the recipient cells to macrophages. Together, our findings reveal a largely unexplored aspect of innate immunity and provide a mechanistic model in which EXi prime local and distant naïve monocytes via p38-induced differentiation and IL-6 production to protect against infection with Gram-negative bacteria.

## Introduction

The role of vesicular trafficking and transport in regulating innate immune responses remains poorly defined. In particular, developing a deep molecular understanding of the contributions of small extracellular vesicles (sEVs) is still at its infancy. As carriers of biological molecules (proteins, mRNAs, micro-RNAs, and lipids), sEVs travel from their site of production to target sites, either within the same microenvironment or to sites distant from their origin, where they bring about physiological changes ^1^. To date, sEVs have been implicated in the progression of a variety of human diseases, such as cancer, neurological disorders, cardiovascular diseases, and infectious diseases ^2–5^. In the context of infection, sEVs have been shown to either activate or inhibit the immune system, depending on the pathogen ^6,7^. As such, sEVs represent promising novel strategies for vaccinations, therapeutics, and diagnostics, since they are produced naturally, can be detected in and purified from bodily fluids, and their contents are disease specific and functionally active in recipient cells. For a few infections, the specific packaging of sEVs in response to infection and the resulting modulation of the immune response have been reported ^8–10^. However, several significant gaps in the knowledge of how sEVs regulate innate immunity still exist, including the regulation of protein signaling in recipient innate immune cells such as monocytes and how this regulation impacts bacterial spread throughout the course of the host response to infection.

There are currently limited investigations into sEV function during infection with highly pathogenic Gram-negative bacterial agents such as *Yersinia pestis* (Yp) and *Burkholderia pseudomallei* (Bp), the causative agents of plague and melioidosis, respectively ^4,11,12^. Given that the virulence factors of both Yp and Bp and their associated host targets are well characterized, these bacteria serve as excellent models for studying Gram-negative bacterial pathogens. Furthermore, reflecting grave concerns over their potential use for bioterrorism, Yp and Bp have been classified as pathogens of high concern (Category A and Category B) ^13,14^. Due to an increasing rise of incidence, plague has also been classified as a re-emerging infectious disease ^11^. However, despite these concerns and challenges, no approved vaccine or highly effective therapeutic interventions currently exist ^15^.

Here, we have used models of infection for Yp and for *Burkholderia thailandensis* (Bt), which is a well-established surrogate for Bp, to investigate the regulatory effects of treating naïve human monocytes with purified sEVs obtained from either Yp-infected cells (EXi-Yp) or Bt-infected cells (EXi-Bt), addressing the significant gap in knowledge of the molecular mechanisms by which sEV function regulates innate immune response to infection with Gram-negative bacteria. We analyzed EXi-induced immune responses in human monocytes and how these changes influence intracellular bacterial growth/survival. As part of this effort, we employed our well-established reverse phase protein microarray platform to comprehensively survey the signaling pathway changes in EXi-treated monocytes, enabling identification of the signaling effects that lead to the observed phenotypes. Using high sensitivity and high specificity mass spectrometry (MS) analysis, we also identified bacterial protein moieties that become selectively packaged in EXi-Yp. To further validate our in vitro findings, we analyzed EXi-Yp exchange effects using our microfluidic chip platform that allows functional interrogation of EVs under physiologically relevant conditions. Furthermore, we tested EXi-Yp protective effects against Yp infection using a well-established mouse model of plague. Collectively, our results suggest a mechanistic model of how EXi-Yp and EXi-Bt can be used as part of the host arsenal to combat infection. In this model, the EXi produced by infected immune cells prime local and distant naïve monocytes to mount an immune response similar to direct infection. This includes the modulation of a defined small set of signaling pathways, including activation of p38 that triggers monocyte to macrophage differentiation and release of specific pro-inflammatory cytokines that includes IL-6. The substantial upregulation of IL-6 release in turn induces significantly increased bacterial clearance, providing protection against infection. These findings make a notable contribution to our developing understanding of the role of sEVs in innate immunity and provide insights into key host response mechanisms that may aid future development of innovative countermeasures against bacterial pathogens.

## Results

### Purification and Characterization of sEVs

For all the studies reported here, we used purified sEVs that were prepared from cell culture supernatants as described previously ^16^ with some slight modifications (Fig. S1). We validated our purification procedure by probing the density gradient fractions for the common sEV markers CD63, TSG101, and Flotillin-1 (Fig. 1a). Consistent with previously published reports ^17^, sEVs obtained from either uninfected or Yp-infected U937 human monocytes migrate to the 1.103 g ml^-1^ and 1.149 g ml^-1^ density fractions (Fig. 1a). ZetaView nanoparticle tracking analysis (NTA) was performed on both EXu and EXi-Yp. The EXu and EXi-Yp preparations had similar concentrations (Fig. 1b), and population mean diameters (Fig. 1c) that are within the accepted sEV size range of less than 200 nm ^17,18^. The former observation indicates that, unlike what has been reported for human polymorphonuclear neutrophils^19^, Yp infection did not impact sEV release from human monocytes^19^. EXu and EXi-Yp also showed similar Zeta potential values (Fig. 1d), indicating high vesicle stability. Furthermore, SEM analysis of purified EXi-Yp revealed the presence of intact vesicles (Fig. 1e).

**Figure 1.**
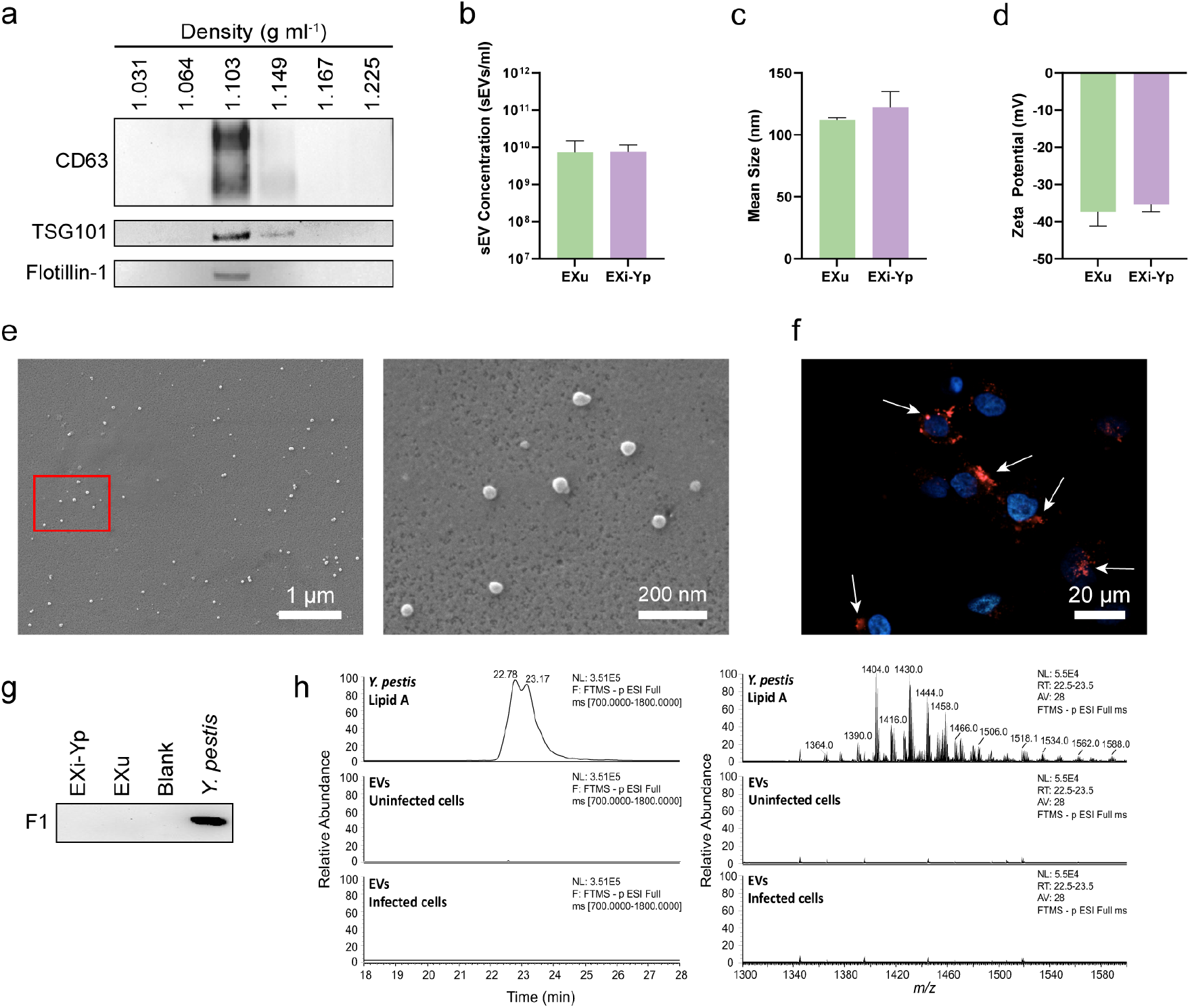
Purification and Characterization of sEVs released by *Y. pestis* infected cells (EXi-Yp). **(a)** Purification of EXi-Yp by density gradient centrifugation and western blot analysis of collected fractions for the presence of sEV markers: CD63 (top), TSG101 (middle), and Flotillin-1 (bottom). **(b)** sEV concentration, **(c)** mean sEV diameter, and **(d)** Zeta potential of purified EXu and EXi-Yp measured by ZataView nanoparticle tracking analysis (mean ± SD, n=3). **(e)** SEM of purified EXi-Yp demonstrates the presence of intact vesicles. A representative image from three biological repeats (n=3) is shown **(f)** Confocal imaging of PKH-stained EXi-Yp (red) inside DAPI stained (blue) recipient cells at 3 h post-treatment. White arrows point to labeled sEVs. **(g)** Western blot analysis of purified EXi-Yp to probe for the presence of Yp F1 protein that serves as an OMV marker. **(h)** MS1 spectra of *Y. pestis* lipid A species detected by t-SIM at 22.5-23.5-min retention time (left) and Targeted selected-ion monitoring (t-SIM) chromatogram of *Y. pestis* lipid A signature parent ion-species (right). Representative plots from three biological replicates are shown (n=3).

To analyze the uptake of purified sEV preparations by recipient monocytes, cells were treated with sEVs labeled with lipophilic PKH dye and imaged by confocal microscopy. The results demonstrated sEV uptake into the recipient cells (Fig. 1f), with a plateau in uptake efficiency observed within 3-4 hours post sEV addition. As bacterial outer membrane vesicles (OMVs) overlap with the size range of host sEVs, we also investigated whether our purified preparations carry OMVs using both western blot analysis and high sensitivity mass spectrometry (LC/MS/MS) studies. Western blot analysis showed a lack of the Yp OMV marker F1 antigen (Fig. 1g). Consistent with this observation, LC-MS/MS analysis failed to detect the presence of the lipid A moiety of Yp LPS ^20^ in purified EXi-YP (Fig. 1h), demonstrating that bacterial OMVs are not detected in sEVs purified from Yp-infected cells.

### EXi-Yp Induce Human Monocyte Differentiation to Macrophages

Previous reports have demonstrated that for some viral infections the EXi released from infected cells affect the viability of naïve recipient cells^21^. We analyzed whether a similar phenomenon is observed in our infection models. Unlike certain viral infections, we did not observe a reduction in the viability of naïve U937 recipients following treatment with EXi-Yp (Fig. S2). However, naïve U937 monocytes treated with EXi-Yp did show a drastic reduction in their rate of growth (about five-fold) over the initial 24 hours post treatment, as compared to untreated cells or cells exposed to a matched amount of EXu (Fig. 2a).

**Figure 2.**
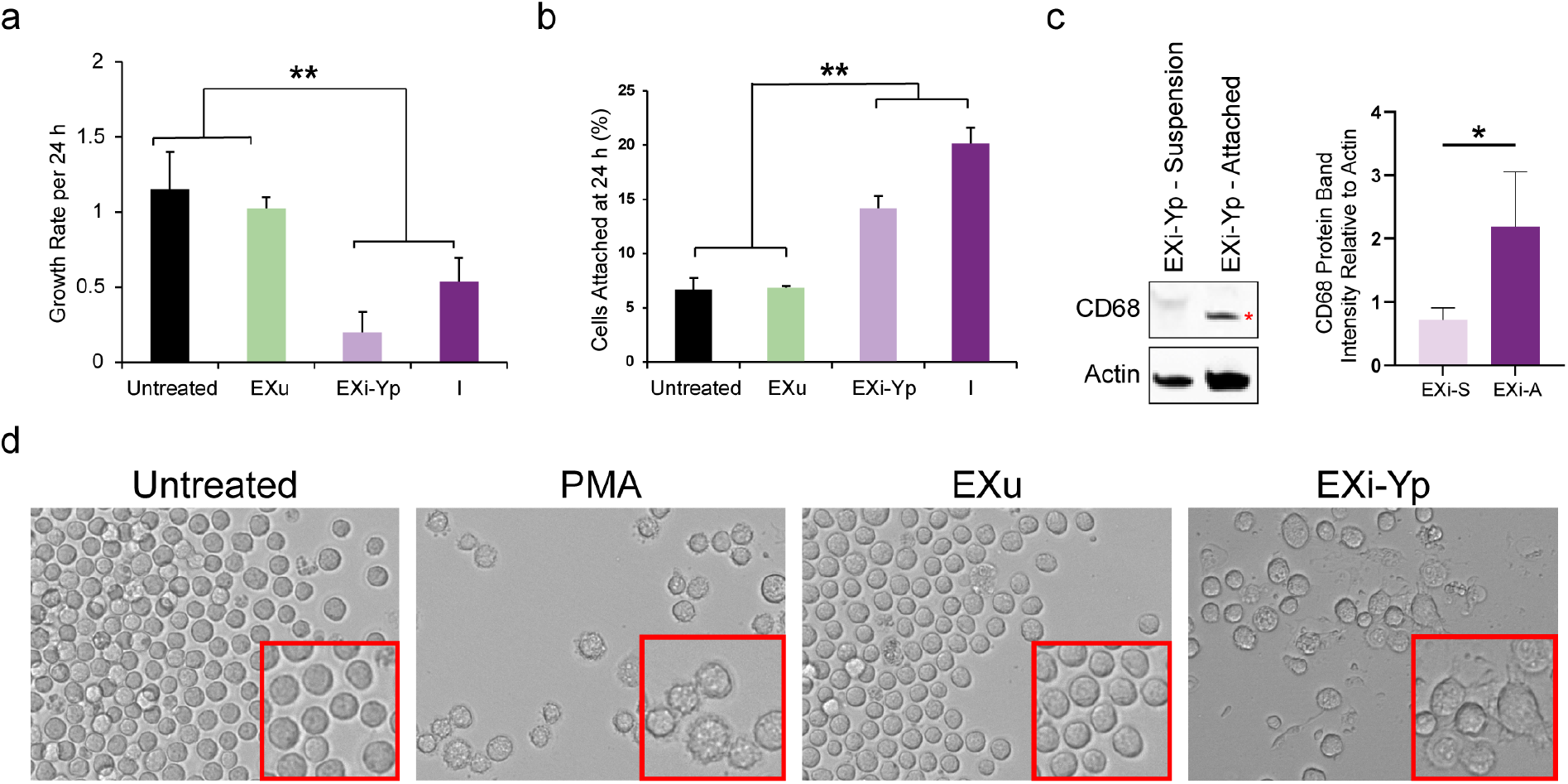
EXi-Yp induce differentiation of naïve human monocytes to macrophages. **(a)** Cell growth rate from 0 h to 24 h of U937 cells that were either left untreated, or were treated with equivalent numbers of EXu or EXi-Yp, or were infected with Yp (means ± SEM, n=5). **(b)** Cell attachments were quantified by trypan blue assay at 24 h post-treatment for U937 cells that were either left untreated, or were treated with equivalent numbers of EXu or EXi-Yp, or were infected with Yp, (means ± SEM, n=3). **(c)** U937 cells were treated with EXi-Yp and incubated for 24 h. Suspension cells (S) and attached cells (A) were isolated and probed by western blot for the macrophage differentiation marker CD68 (marked by red star). Band intensities were quantified relative to actin (mean ± SD, n=3). **(d)** Morphological changes of naïve U937 cells left untreated or treated with PMA, EXu, or EXi-Yp observed using brightfield microscopy at 96 h post-treatment.

This response was similar to the cells infected with Yp bacteria (Fig. 2a). For either the EXi-treated cells or Yp-infected cells, this effect is not observed past the initial 24 hours post treatment (Fig. S3a and S3b). Furthermore, this differential growth response is a unique consequence of EXi-Yp treatment of host cells, as no direct effect on Yp survival/growth is observed when Yp cells are directly treated with EXi-Yp (Fig. S3c). A retardation in the growth rate of monocytes, including in the growth rate of U937 cells as a result of G1 arrest, is one hallmark of monocyte to macrophage differentiation ^22^.

Consequently, we next analyzed whether EXi-treated cells show increased attachment, another hallmark of the macrophage differentiation phenotype ^23^. Treatment of U937 monocytes with EXi-Yp induced a significant increase in cell attachment (14 ± 1%) as compared to either EXu-treated or untreated cells (6 ± 0.1% and 6 ± 1% respectively) (Fig. 2b). Also, as with the cell growth studies, a similar increased attachment phenotype was observed for cells infected with Yp, with 20.1 ± 1% of the infected cells showing attachment. To further verify EXi induction of differentiation, we also tested for increased expression of the macrophage differentiation marker CD68 ^24^ in cells that attached in response to treatment with EXi-Yp. Consistent with the observed growth rate and attachment phenotypes, cells treated with EXi-YP showed a clear increase in CD68 expression within the attached subpopulation when compared to cells remaining in suspension (Fig. 2c). Finally, we treated U937 monocytes with either PMA, a known stimulus of differentiation ^24^, or with equivalent amounts of EXu or EXi-Yp, and visually compared them to untreated cells (Fig. 2d). While no differences were observed between the untreated and EXu-treated conditions, both EXi-treated and PMA-treated cells showed notable changes in comparison, exhibiting protruding membranes and also flatter, spread-out morphologies that result from increased surface adherence. Collectively, these results demonstrate that treatment with EXi-Yp triggers the differentiation of human monocytes to macrophages, an innate immune response that mimics the autocrine immune reaction of cells to direct infection with Yp.

### EXi-Yp Treatment Induces Pro-inflammatory Cytokine Release and IL-6-dependent Bacterial Clearance

Having observed the differentiation phenotype, we next analyzed whether EXi-Yp treatment also elicits cytokine release, another facet of innate immune activation. A 10-plex cytokine array panel was used to measure the levels of the human pro-inflammatory cytokines IFNγ, IL-1α, IL-1β, IL-2, IL-4, IL-6, IL-8, IL-10, IL-12p70, and TNFα. Compared to untreated and uninfected cells, or cells treated with an equivalent amount of EXu, treatment with EXi-Yp resulted in significantly increased release (10- to 100-fold) of IL-6, IL-8, and IL-10 from 12 hours to 48 hours post treatment (Fig. 3a), reminiscent of the reported cases of co-regulation of these three cytokines ^25,26^. These cytokines were also induced when the cells were infected with Yp, demonstrating further similarity between host response to EXi-Yp treatment and infection with Yp. Given the EXi-mediated induction of cytokine release and monocyte-to-macrophage differentiation, we analyzed whether treatment with EXi-Yp has an effect on bacterial clearance in U937 cells that have undergone EXi-induced differentiation. U937 cells were pretreated with equivalent amounts of either EXi-Yp or EXu for 24 hours, and subsequently the suspension cells were removed and the remaining attached cells were infected with Yp to quantify both the amount of Yp uptake and Yp survival/growth during 24 hours post-infection. Our results show that pre-treatment with EXi-Yp, but not EXu, significantly increased the amount of Yp clearance by the host cells over the period of 24 hours post-infection (Fig. 3b).

**Figure 3.**
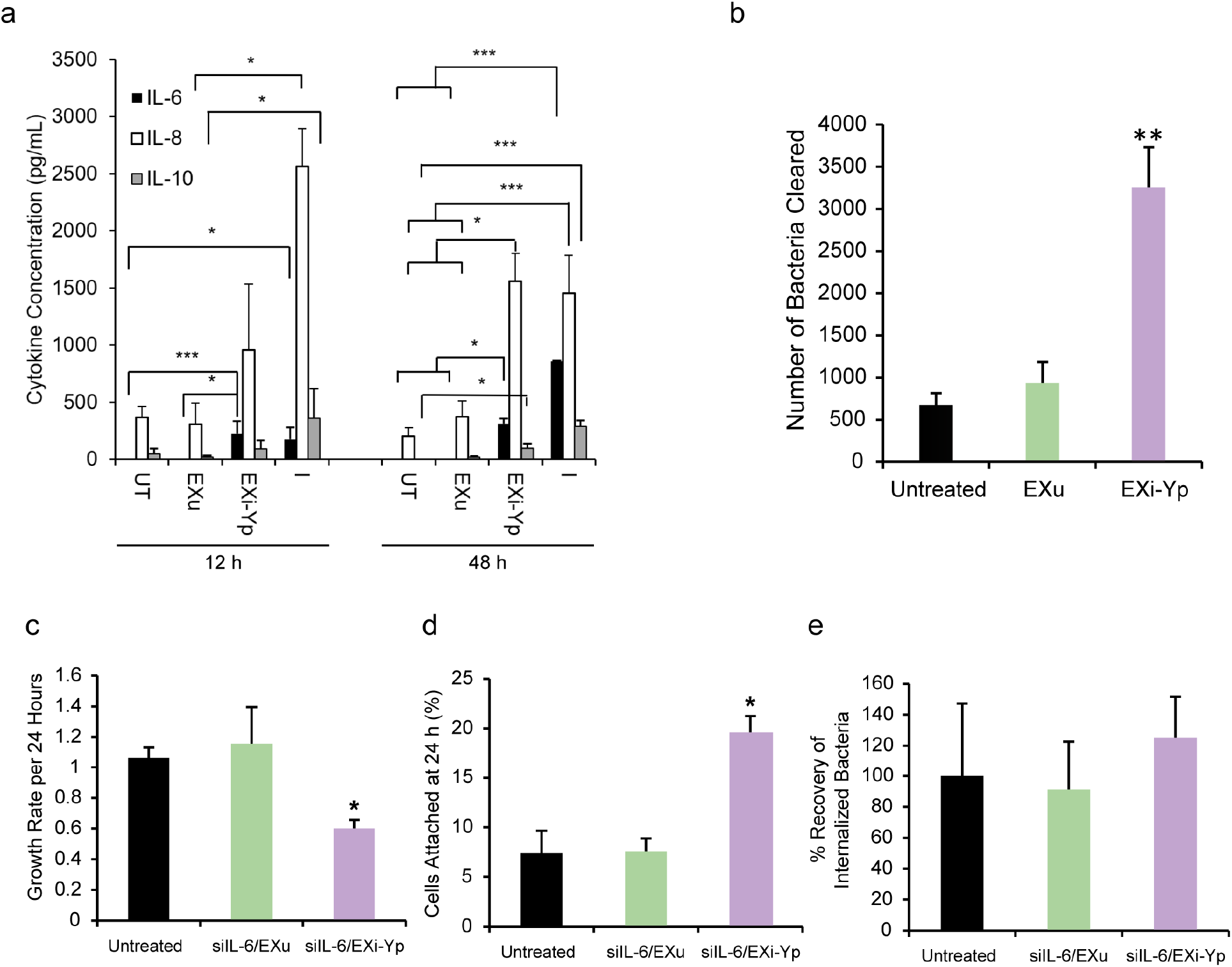
EXi-Yp treatment induces pro-inflammatory cytokine release and IL-6 dependent bacterial clearance. **(a)** U937 cells were either left untreated, or were treated with equivalent numbers of EXu or EXi-Yp, or were infected with Yp. Culture supernatants were harvested at 12 h and 48 h post-treatment for quantitative multiplex probing of a panel of 10 pro-inflammatory cytokines. Measurements for cytokines that were significantly induced by EXi-Yp are presented (mean ± SEM, n=4). **(b)** Naïve U937 cells that were either left untreated or were pre-treated with equivalent numbers of EXi-Yp or EXu for 24 h, were infected with Yp. Internalized bacteria were recovered and quantified by CFU counts at 24 h post-infection (mean ± SEM, n=4). **(c)**, **(d)**, and **(e)** U937 cells were either left untreated or were treated with siIL-6 siRNA or scrambled control siRNA for 24 h and subsequently incubated with EXu or EXi-Yp for 24 h: **(c)** growth rate from 0 h to 24 h post-treatment (mean ± SEM, n=3); **(d)** cell attachment at 24 h (mean ± SEM, n=3); and, **(e)** quantitation of bacterial clearance at 24 h post-Yp infection (mean ± SEM, n=3).

We examined whether IL-6 or IL-8 release influences any of the observed EXi-induced effects. For these experiments, we silenced IL-6 or IL-8 in recipient naïve monocytes using IL-6 siRNA or IL-8 siRNA respectively (Fig. S4a), followed by treatment with EXi-Yp and analysis of EXi-induced responses. Silencing of IL-8 had no effect on the observed phenotypes. However, while Silencing of IL-6 in recipient cells showed no effect on either the EXi-induced cell growth defect (Fig. 3c) or the EXi-induced cell attachment phenotype (Fig. 3d), it resulted in a significant reduction of EXi-induced bacterial clearance (Fig. 3e), bringing the observed significant increase in bacterial clearance down to control levels. In contrast, silencing of IL-8 did not result in any changes to the bacterial clearance phenotype (Fig. S4b). These results demonstrate that while increased IL-6 release in response to EXi-Yp treatment does not affect the differentiation phenotype, it is required for increased bacterial clearance.

### A Distinct Small Set of Signaling Pathways are Modulated by EXi-Yp

To address the significant gap of knowledge in how cellular pathways are modulated by sEVs from bacterially infected cells, we probed the protein pathway responses to EXi-Yp at multiple timepoints post treatment using two fully independent sample preparations. For this analysis, we used 173 pre-validated and highly specific antibodies to probe for numerous signaling pathways across the cell, including those that are known to be involved in the host response to infection (Table S1). We used our previously established and highly validated quantitative RPPA platform ^27–29^ to conduct these high throughput studies, analyzing all 173 target proteins across all treatment conditions and at multiple timepoints post treatment (Fig. 4).

**Figure 4.**
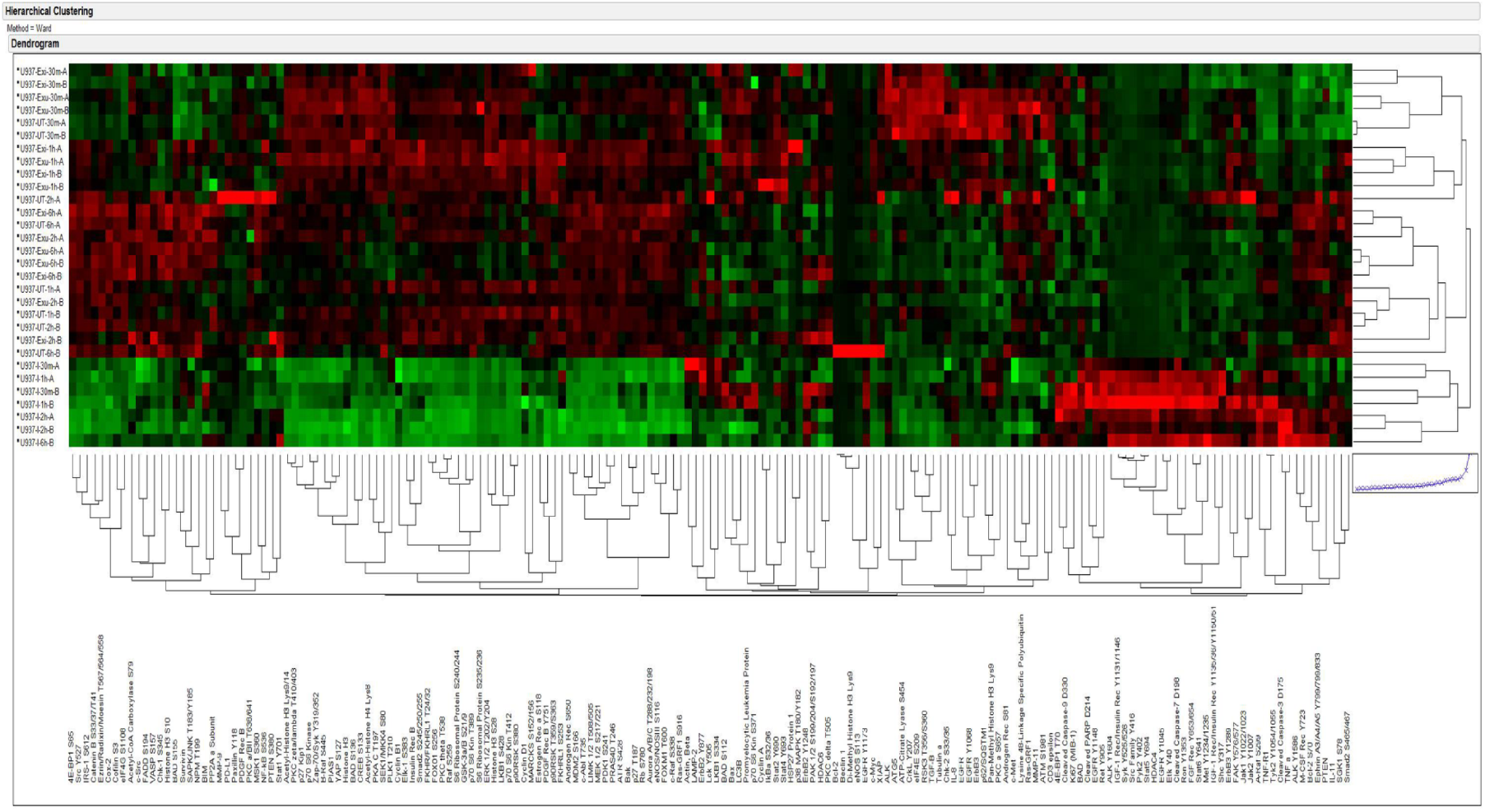
Signaling pathway modulation following EXi-Yp treatment. Naïve U937 cells were either left untreated, or were treated with equivalent numbers of EXi or EXu, or were infected with Yp. At 30 min, 1 h, 2 h, and 6 h post-treatment, whole cell lysates were prepared and probed using 173 validated antibodies to quantitatively survey changes across cell signaling pathways. The heat map shows unsupervised hierarchical 2-way clustering analysis of all the samples. Rows represent different samples, and columns show antibodies tested. Relative signal intensity was assigned based on the lowest signal on the array and depicted by color intensity, with red color representing higher signal and green color representing lower signal.

Analysis of the cells treated with EXi-Yp demonstrated a strong modulation of a distinct set of cell signaling molecules when compared to control conditions (Table 1). Consistent with EXi-induced inhibition of cell growth, at 30 minutes post-treatment, we observed both a significant increase in total levels of the BAD protein, a potent pro-apoptotic activator that inhibits cell growth, and a significant decrease in the activation of the pro-cell growth signaling protein FAK. Furthermore, we observed a decrease in IGF-1 Rec activation. This finding is consistent with the observed up-regulation of total BAD levels given that IGF-1 signaling negatively regulates BAD through its activation of AKT ^29,30^. Interestingly, at 2 hours post treatment, we observed a significant up-regulation of p38 MAPK activation, as well as significant inhibition of the pro-growth cell cycle regulators ALK and JAK2. Because activation of p38 MAPK has been shown to cause all the EXi-Yp-induced phenotypes reported here, our RPPA findings pointed to p38 as a primary key regulator of the EXi-Yp-associated responses. In addition, it is noteworthy that inactivation of JAK2 has been shown to cause both G1 cell cycle arrest ^31^ and macrophage differentiation ^32^, and the inactivation of ALK inhibits G1/S cell cycle progression ^33^.

**Table 1.** RPPA Identification of Significant Signaling Changes in EXi-Yp Treated Cells as Compared with Untreated Cells and EXu-treated Condition.

| 30 min Post-Treatment | Bacterial Uptake | Growth/Cell Cycle | Differentiation | Cytokine Release |
| --- | --- | --- | --- | --- |
| BAD |  | Expression inhibits cell growth |  |  |
| Pan-Methyl Histone H3 Lys9 |  |  | Deactivated during astrocyte differentiation |  |
| FAK Y576/577 |  | Activation critical for G1/S cell cycle progression |  |  |
| IGF-1 Rec/Insulin Rec Y1135/36/Y1150/51 |  | Activation stimulates growth |  |  |
| 2 h Post-Treatment | Bacterial Uptake | Growth/Cell Cycle | Differentiation | Cytokine Release |
| Acetyl-CoA Carboxylase S79 |  | Inhibition decreases cell growth |  |  |
| ALK Y1604 |  | Activation induces G1/S cell cycle progression | Activation induces neuronal differentiation |  |
| ERM T567/564/558 |  | Active form transports growth signals |  |  |
| p38 MAPK T180/Y182 | Active form induces phagocytosis | Active form induces G1/S arrest | Active form induces M2 differentiation | Active form can induce IL-6, IL-8 and IL-10 release |
| Jak2 Y1007 |  | Inactivation causes G1 cell cycle arrest | Deactivated during macrophage differentiation |  |

### Increased p38 Activity Is a Main Regulator of EXi-Yp-induced Phenotypes

Given the reported association of p38 activity with the phenotypes that are induced by the EXi, we assessed how inhibition of p38 activity in U937 cells affects the EXi-induced phenotypes. To achieve p38 inhibition, we made use of the p38 inhibitor PH-797804, which has been shown to inhibit p38α-induced phosphorylation in HSP-27 in U937 cells ^34^. We first determined the dosage and treatment duration at which p38-induced HSP-27 phosphorylation is inhibited without any adverse effects on cell viability (Fig. S5a and b). Using these parameters, we observed that inhibition of p38 activity with PH-797804 led to a significant reversal of EXi-Yp-induced growth, attachment, and IL-6 production phenotypes (Fig. 5a-c).

**Figure 5.**
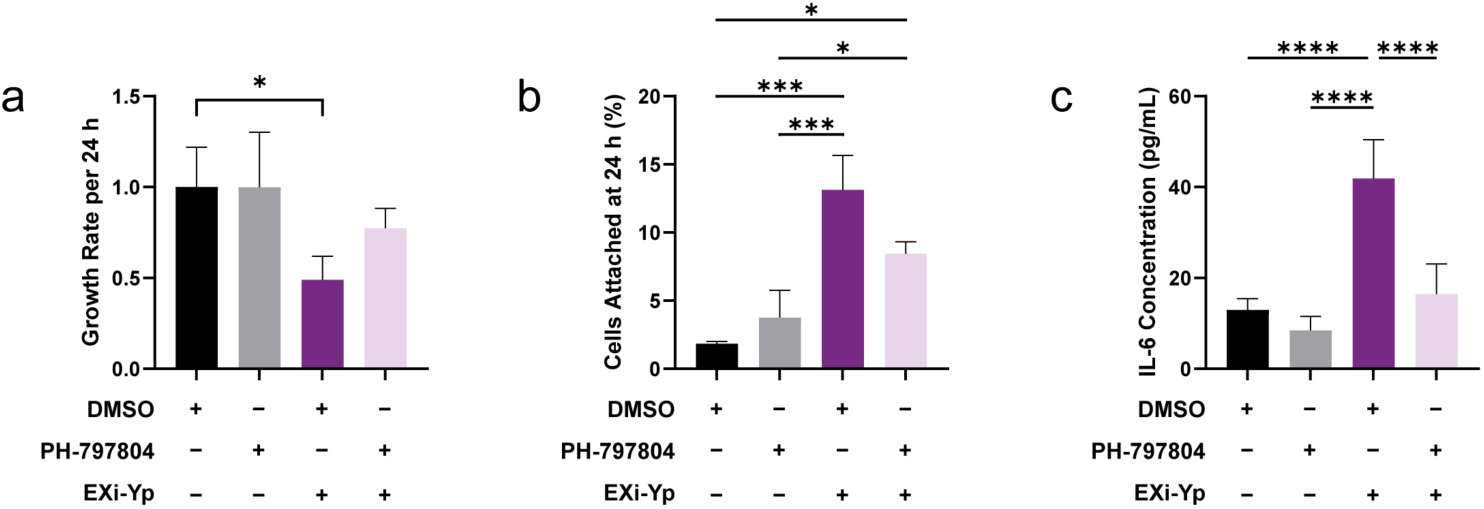
Increased p38 signaling regulates innate immunity effects of EXi-Yp. U937 cells pre-treated with p38 inhibitor or DMSO were either left without any further treatments or were treated with EXi-Yp: **(a) g**rowth rate from 0 h to 24 h post treatment (mean ± SD, n=3); **(b)** cell attachment at 24 h post treatment (mean ± SD, n=3); and, **(c)** IL-6 concentration of cell culture supernatant at 48 h post treatment (mean ± SD, n=4).

### sEV Protein Components Contribute to EXi-Yp Functional Effects

To analyze the potential role of sEV surface proteins in the activation of host innate immunity, we treated purified EXi with the general protease proteinase K to remove external protein moieties. To verify that our treatment conditions do not compromise the structural integrity of purified sEVs, proteinase K-treated EXi-Yp (PK-EXi-Yp) were analyzed by TEM and confirmed to have remained as intact vesicles (Fig. S6a). We next analyzed whether the removal of external protein moieties associated with EXi-Yp affects internalization into recipient cells. Comparative flow cytometry analysis of naïve monocytes treated with fluorescently tagged EXu, EXi-Yp, and PK-EXi-Yp, demonstrated lower uptake levels for the proteinase K-treated sample, although not to the extent that was found to be statistically significant (Figure S6b). We found that PK-EXi failed to either significantly reduce growth rate or increase cell attachment (Fig. S6c and d), and also failed to significantly increase bacterial uptake (Fig. S6e). These PK-treated vesicles also failed to induce significant induction of Il-6, IL-8, and IL-10; in particular, IL-6 production reverted back to basal levels (Fig. S6f). These results demonstrate that the EXi-Yp induction of differentiation and IL-6-dependent increase in bacterial clearance all require engagement of sEV surface protein(s).

We have also performed LC-MS/MS analysis of four biological replicates of EXi-Yp to determine whether they carry any bacterial proteins. For each replicate, equivalent numbers of purified EXu were also analyzed in parallel as control. We identified three specific Yp proteins in all 4 independently purified EXi-Yp samples, indicating that they are specifically packaged into EXi-Yp during infection: i) molecular chaperone GroEL, ii) elongation factor Tu, and iii) subunits of the Urease enzyme (Table 2). Of particular note is that all three proteins have been shown to be antigenic during other bacterial infections^35–41^.

**Table 2.** *Yersinia pestis* proteins associated with EXi-Yp.

| EXi-Associated Yp Proteins | Function | Antigenic Properties |
| --- | --- | --- |
| Urease Subunits $\alpha$ and $\gamma$ | Catalyzes the breakdown of urea <sup>42</sup> | Antigenic in *Yersinia enterocolitica* and *Helicobacter pylori* <sup>35,43</sup> |
| GroEL | Molecular chaperone required for bacterial growth <sup>44</sup> | Antigenic in *Burkholderia pseudomallei* and *Mycoplasma gallisepticum* <sup>37,38</sup> |
| Elongation factor Tu | Involved in protein synthesis <sup>45</sup> | Antigenic in *Burkholderia pseudomallei* and *Borrelia burgdorferi* <sup>39,40</sup> |

Similar to many other Gram-negative bacteria, Yp possesses a type III secretion system (T3SS), which is used to deliver various cytotoxic effector proteins, including *Yersinia* outer-membrane proteins (Yops), into the host cell ^46^. Functions of these Yops include modulation of apoptosis, phagocytosis, and cytokine production ^47^. Given the importance of this secretion system in immune evasion and progression of Yp infection, we assessed whether the T3SS of Yp influences the biogenesis, and consequently the functional properties, of the EXi-Yp. The EXi were purified from the culture supernatant of U937 cells infected with a Yp strain that lacks the pCD1 plasmid, and thus the T3SS ^48^. We demonstrate that these sEVs (designated as EXi-YpΔT3SS) are still capable of inducing reduced cell growth rate and increased cell attachment phenotypes in recipient naïve U937 cells (Fig. S7a and b). In contrast, while EXi-YpΔT3SS still induces increased release of IL-6 in recipient cells, the level of increase is considerably lower than that observed in cells treated with EXi-Yp (compare Fig. S7c with the 48 hr time point in Fig. 3a). These results suggest that components of the T3SS may be important for the ability of EXi-Yp to induce IL-6, which is required for the bacterial clearance phenotype as shown in Fig. 3e.

### Purification and Characterization of sEVs from Bt-Infected Cells

To determine whether the EXi-Yp effects are exclusive to YP or rather represent common mechanisms by which host sEVs regulate innate immune responses to Gram-negative pathogens, we also analyzed the Bt infection model. For these studies, using the same purification process that was used for EXi-Yp, we purified sEVs from U937 cells infected with Bt (EXi-Bt), which is closely related to Bp and serves as a well-accepted model for Bp studies at the BSL-2 level ^49^. Similar to the results for EXi-Yp and consistent with reported sEV buoyant density range, sEVs obtained from either uninfected or Bt-infected U937 human monocytes migrated to 1.103 g ml-1 and 1.149 g ml-1 density fractions (Fig. 6a), and probing of the density gradient fractions by western blot demonstrated the presence of the CD63, TSG101, and Flotillin-1 markers in the sEV fractions (Fig. 6a). ZetaView particle tracking analysis demonstrated that Bt infection does not significantly impact the quantity of recovered sEVs, similar to our observation for Yp infection (Fig. 6b). It also showed that, similar to EXu, the EXi-Bt mean diameter is within the accepted sEV size range (Fig. 6c), with Zeta potential value indicating high vesicle stability (Fig. 6d). TEM analysis of purified sEVs demonstrated that our isolation process resulted in intact vesicles (Fig. 6e).

**Figure 6.**
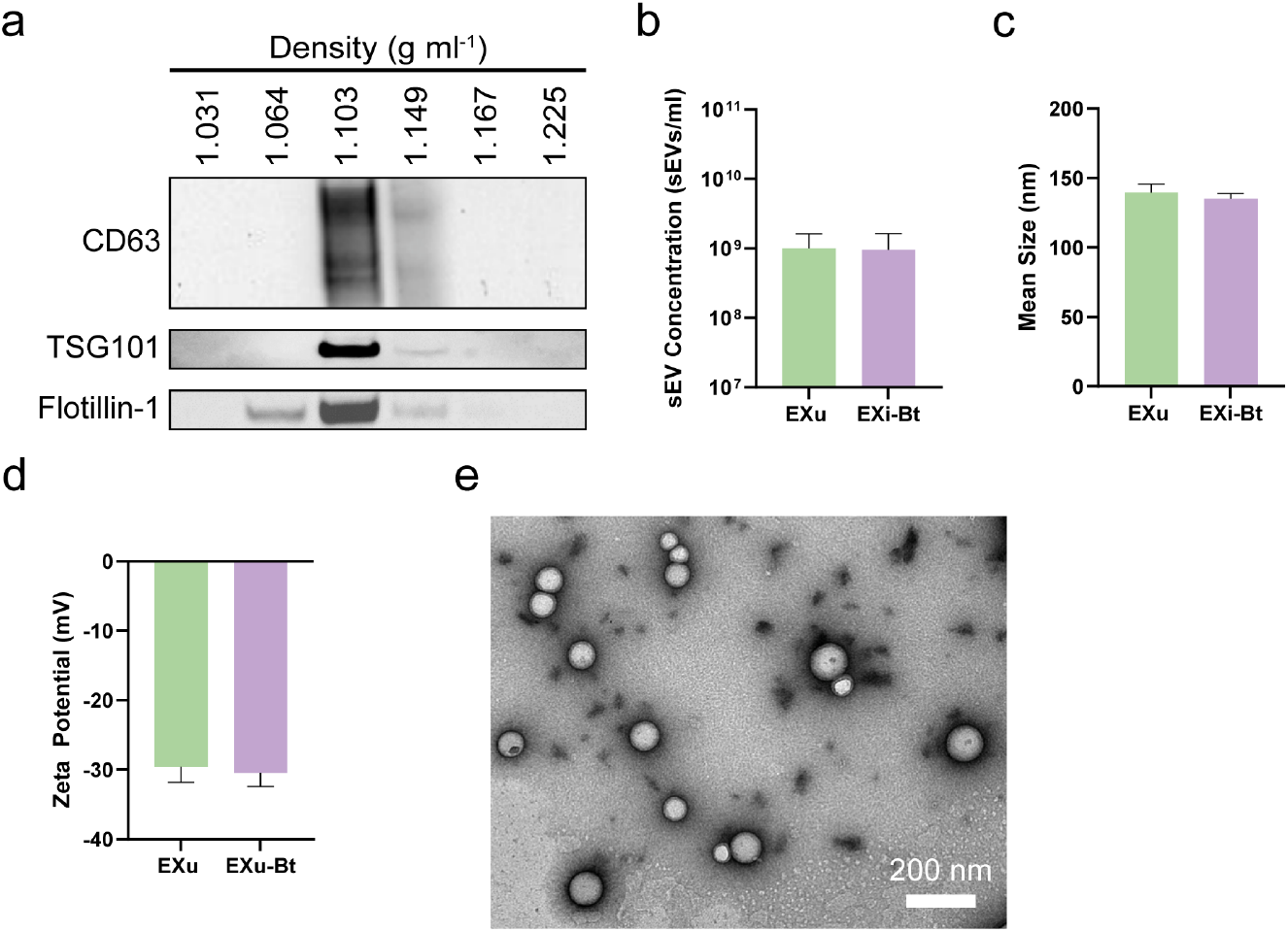
Purification and Characterization of sEVs released by *B. thailandensis* infected cells (EXi-Bt). **(a)** Purification of EXi-Bt by density gradient centrifugation and western blot analysis of collected fractions for the presence of sEV markers: CD63 (top), TSG101 (middle), and Flotillin-1 (bottom). **(b)** sEV concentration, **(c)** mean sEV diameter, and **(d)** zeta potential of purified EXu and EXi-Bt measured by ZataView nanoparticle tracking analysis (mean ± SD, n=3). **(e)** TEM of purified EXi-Bt demonstrates the presence of intact vesicles.

### EXi-Bt Induce Human Monocyte Differentiation to Macrophages

Similar to our EXi-Yp studies, we confirmed that treatment of naïve U937 cells with EXi-Bt did not impact cell viability up to 48 hours post-treatment (Fig. S8). However, as we observed with EXi-Yp, the treatment of naïve U937 monocytes with EXi-Bt showed significantly reduced growth rate (about 2.5-fold) within the first 24 hours post-treatment when compared to untreated cells or EXu-treated cells (Fig. 7a). A similar reduction in cell growth was also observed in cells infected with Bt (Fig. 7a).

**Figure 7.**
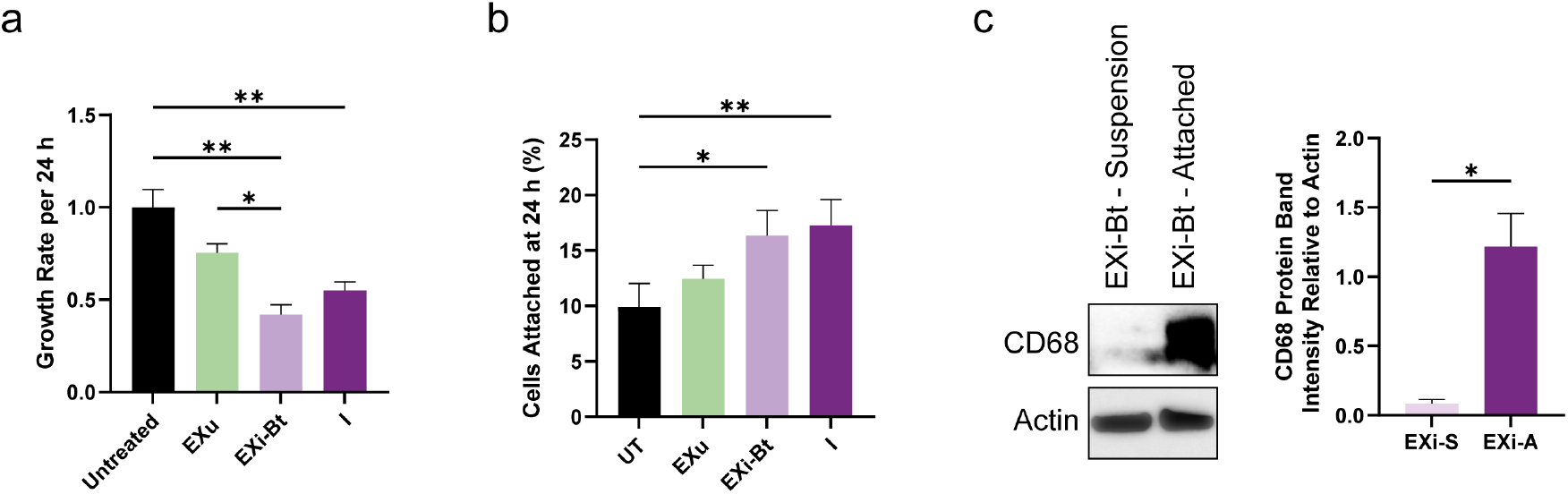
EXi-Bt induce differentiation of naïve human monocytes to macrophages. **(a)** Cell growth rate from 0 h to 24 h of U937 cells that were either left untreated, or were treated with equivalent numbers of EXu or EXi-Bt, or were infected with Yp (mean ± SD, n=3). **(b)** Cell attachments were quantified by trypan blue assay at 24 h post-treatment for U937 cells that were either left untreated, or were treated with equivalent numbers of EXu or EXi-Bt, or were infected with Bt (mean ± SD, n=3). **(d)** U937 cells were treated with EXi-Bt and incubated for 24 h. Suspension cells (S) and attached cells (A) were isolated and probed by western blot for the macrophage differentiation marker CD68. Band intensities were quantified relative to actin (mean ± SD, n=2).

Furthermore, similar to our results with EXi-Yp, treatment of naïve U937 cells with EXi-Bt led to a significant increase in the level of cellular attachment, showing an average attachment rate of 16% (Fig. 7b). A similar level of significant attachment increase was also observed in the Bt-infected U937 cells, showing an average attachment rate of 17% (Fig. 7b). Consistent with these observations, the attached population of EXi-Bt treated cells showed a significant increase in CD68 expression when compared to the population that remained in suspension (Fig. 7c). These results demonstrate that, similar to EXi-Yp, treatment with EXi-Bt induces differentiation of human monocytes to macrophages, mimicking the phenotype that is observed when monocytes are infected with the bacteria.

### EXi-Bt Induce Pro-inflammatory Cytokine Release and IL-6-dependent Bacterial Clearance

We next analyzed whether EXi-Bt can also elicit proinflammatory cytokine release and induce IL-6-dependent bacterial clearance, similar to our observations for EXi-Yp. Using the same 10-plex cytokine array that we used for the EXi-Yp studies, we observed that compared to untreated and EXu-treated cells, treatment with EXi-Bt resulted in significantly increased release (2- to 4-fold) of IL-2, IL-6, and TNF-α at 24 hours post treatment (Fig. 8a). The same set of cytokines were also released at significantly increased levels when the cells were infected with Bt (Fig. 8a), again demonstrating similarities between host responses to EXi-Bt treatment and becoming infected with Bt bacteria. Of particular note is the observation of EXi-Bt induction of IL-6, which as we have shown leads to increased bacterial clearance by EXi-Yp. Therefore, we next analyzed whether treatment with EXi-Bt also leads to increased bacterial clearance in human monocytes. Our results demonstrate that pre-treatment with EXi-Bt, but not EXu, significantly reduced intracellular Bt survival/growth within the host cells over the period of 24 hours post infection (Fig. 8b), mirroring the effects observed with EXi-Yp.

**Figure 8.**
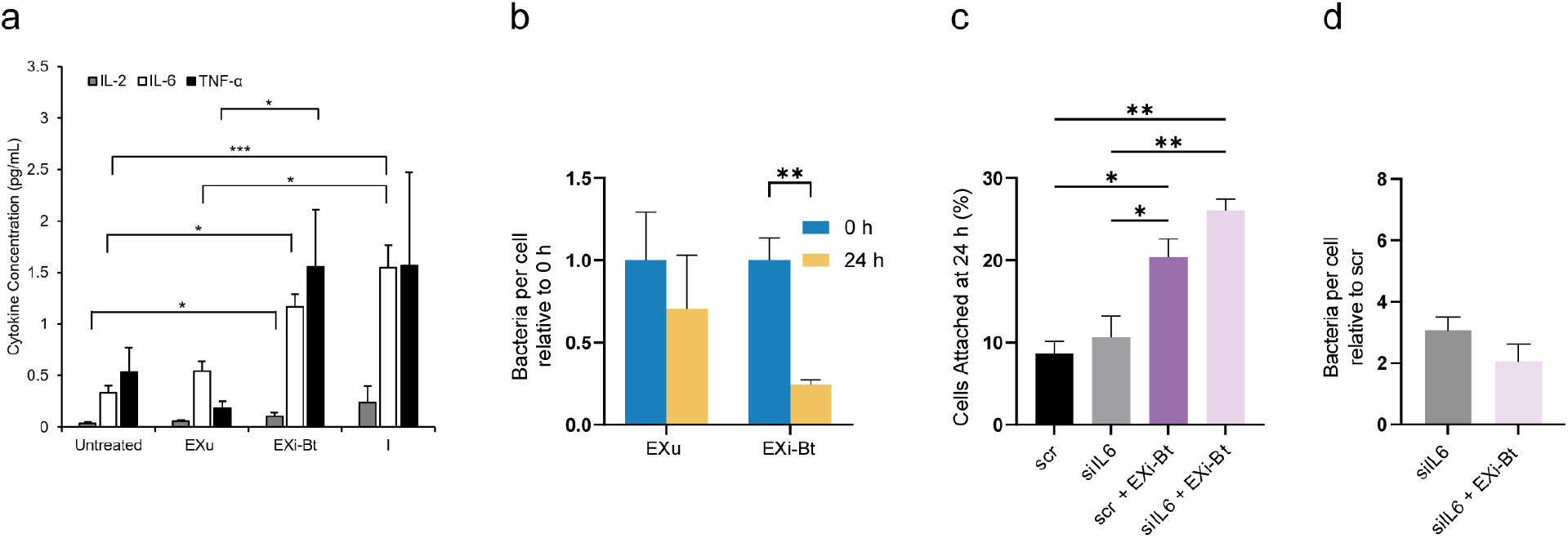
EXi-Bt treatment induces pro-inflammatory cytokine release and IL-6 dependent bacterial clearance. **(a)** U937 cells were either left untreated, or were treated with equivalent numbers of EXu or EXi-Bt, or were infected with Bt. Culture supernatants were harvested at 24 h post-treatment for quantitative multiplex probing of a panel of 10 pro-inflammatory cytokines. The levels of cytokines significantly induced by EXi-Bt are shown (mean ± SEM, n=3). **(b)** Naïve U937 cells that were either left untreated or were pre-treated with equivalent numbers of EXi-Bt or EXu for 24 h, were infected with Bt. Internalized bacteria were recovered and quantified by CFU counts at 0 h and 24 h post-infection (mean ± SD, n=3). **(c)** U937 cells were treated with either siIL-6 siRNA or scrambled control siRNA for 24 h. Subsequently, the cells were incubated with or without EXi-Bt for 24 h and the degree of cell attachment was measured (mean ± SD, n=2). **(d)** U937 cells were treated with either siIL-6 siRNA or scrambled control siRNA for 24 h and subsequently incubated with or without EXi-Bt for 24 h. The attached cell population was then infected with Bt for 24 h, followed by cell lysis and quantification of intracellular bacteria (mean ± SD, n=3).

Based on our findings with EXi-Yp, we next examined whether IL-6 release influences any of the observed EXi-Bt-induced effects. Silencing of IL-6 in recipient cells did not affect the EXi-induced cell attachment phenotype (Fig. 8c). We found that silencing of IL-6 led to increased bacterial burden at 24 hours post infection of U937 cells (Fig. S9), indicating that IL-6 signaling is important for the ability of the cells to control the intracellular growth of Bt. Silencing of IL-6 also resulted in reversal of the EXi-induced decrease in intracellular bacterial burden, with untreated cells and EXi-Bt-treated cells showing similar numbers of bacteria per cell at 24 hours post infection (Fig. 8d). These results show that similar to EXi-Yp, the EXi-Bt population induces significantly increased bacterial clearance that is dependent on IL-6 function, further demonstrating close mechanistic similarities in sEV-dependent host response duirng infection with Gram-negative bacteria.

### Increased p38 Activity Is a Main Regulator of EXi-Bt-induced Phenotypes

As we demonstrated earlier, p38 activity associated with EXi-Yp function is linked to the EXi-Yp-induced phenotypes. Accordingly, we investigated whether increased p38 activity also occurs in response to EXi-Bt, and if so whether it also regulates the same main phenotypes that are induced by the EXi-Bt vesicles (differentiation and IL-6-dependent bacterial clearance). We observed that EXi-Bt treatment induces a significantly increased level of p38 phosphorylation as early as 4 hours post treatment when compared to either untreated cells or EXu-treated cells (Fig. 9a-b). EXi-Bt treatment also leads to slightly increased levels of total p38 (Fig. 9a and c). We also observed that similar to EXi-Yp, inhibition of p38 activity led to a reversal of the EXi-Bt-induced differentiation, as measured by growth rate, cell attachment, and CD68 expression analyses (Fig. 9d and e, Fig. S10). Furthermore, similar to our findings for EXi-Yp effects, p38 inhibition led to a complete loss of the EXi-Bt-induced increase in IL-6 release (Fig. 9f).

**Figure 9.**
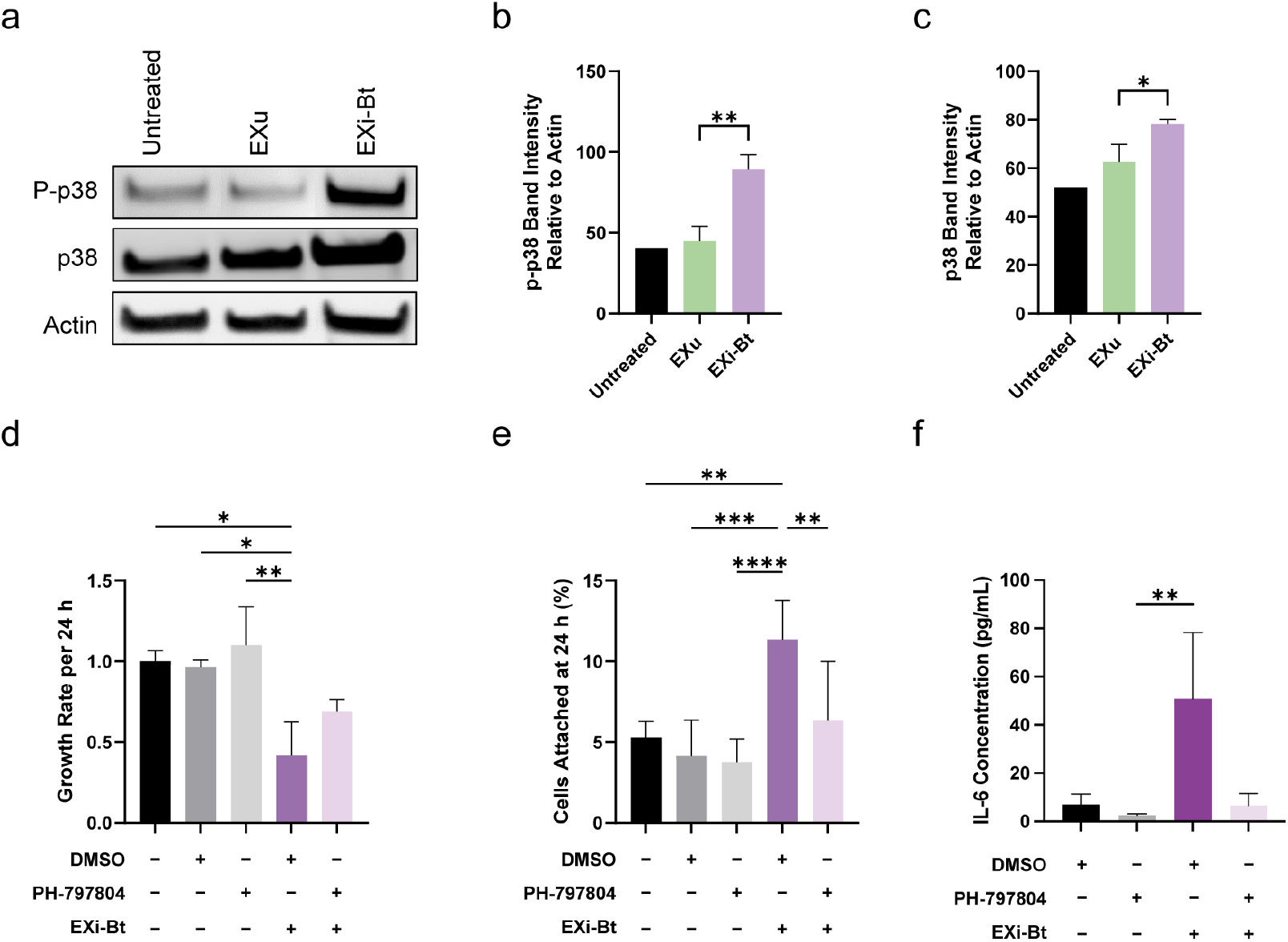
Increased p38 signaling regulates innate immunity effects of EXi-Bt. U937 cells were either left untreated or were treated with equivalent numbers of EXu or EXi-Bt. **(a)** At 4 h post –treatment, cells were lysed and the cell lysates probed for phospho-p38 (p-p38), total p38, and actin by western blot analysis. Band intensities of **(b)** p-p38 and **(c)** p38 were quantified relative to actin (mean ± SD, n=3). **(d)**, **(e)**, and **(f)** U937 cells pre-treated with p38 inhibitor or DMSO were either left without any further treatments or were treated with EXi-Bt: **(d) g**rowth rate from 0 h to 24 h post treatment (mean ± SD, n=3), **(e)** cell attachment at 24 h post treatment (mean ± SD, n=4), and **(f)** IL-6 concentration of cell culture supernatant at 24 h post treatment was measured by ELISA assay for IL-6 (mean ± SD, n=3).

### Intercellular Exchange of EXi-Yp under Physiologically Relevant Conditions Induces Monocyte to Macrophage Differentiation

To provide support for our in vitro findings, we used the microfluidic chip platform that we have developed to allow functional interrogation of EV exchange under more physiologically relevant conditions ^50^. These studies used our platform’s next-generation version that features improved capabilities ^51^. In this platform, intercellular exchange of EVs takes place between a donor cell channel and a recipient cell channel, with the two channels separated by a hydrogel barrier such as the extracellular matrix (ECM)-mimicking Matrigel, which is used to physically separate cell populations confined within microchannels, and mimics tissue environments to enable direct study of EV function (Fig. 10a). To test whether EXi-Yp can induce macrophage differentiation of naive monocytes injected into the recipient channel, we placed EXi-Yp in the donor channel of the chip and analyzed the effects on recipient naive monocytes after 48 hours, allowing sufficient time for efficient translocation of the sEVs across the Matrigel for interaction with the recipient monocytes. A chip with an equivalent number of EXu placed in the donor channel, and also a chip with PMA injected into the donor channel, were also analyzed side by side. The results showed that EXi-YP transmigration across the chip and transfer to recipient monocytes induced their differentiation to macrophages, as indicated by increased cell attachment with flatter, spread-out morphology (Fig. 10b, red arrows). The same effect was observed on the positive-control PMA chip, whereas the negative-control EXu chip did not show any effects (Fig. 10b, yellow arrows). In another set of experiments, we placed Yp-infected cells in the recipient channel of a Matrigel chip and allowed for intercellular exchange of released EXi-Yp with naive recipient monocytes. The results showed that this real-time cell-cell exchange, which represents a more physiologically relevant state, also results in macrophage differentiation of monocytes in the recipient channel (Fig. 10d). To further validate that intercellular sEV exchange is responsible for the observed differentiation phenotype, we also tested a chip with PEGDA-10K as the hydrogel barrier, which has a pore size of about 50 nm and prevents passage of sEVs. Our results showed an absence of the differentiation phenotype on the PEGDA-10K chip (Fig. 10d), supporting the conclusion from our earlier *in vitro* studies that EXi-Yp vesicles induce monocyte to macrophage differentiation.

**Fig 10.**
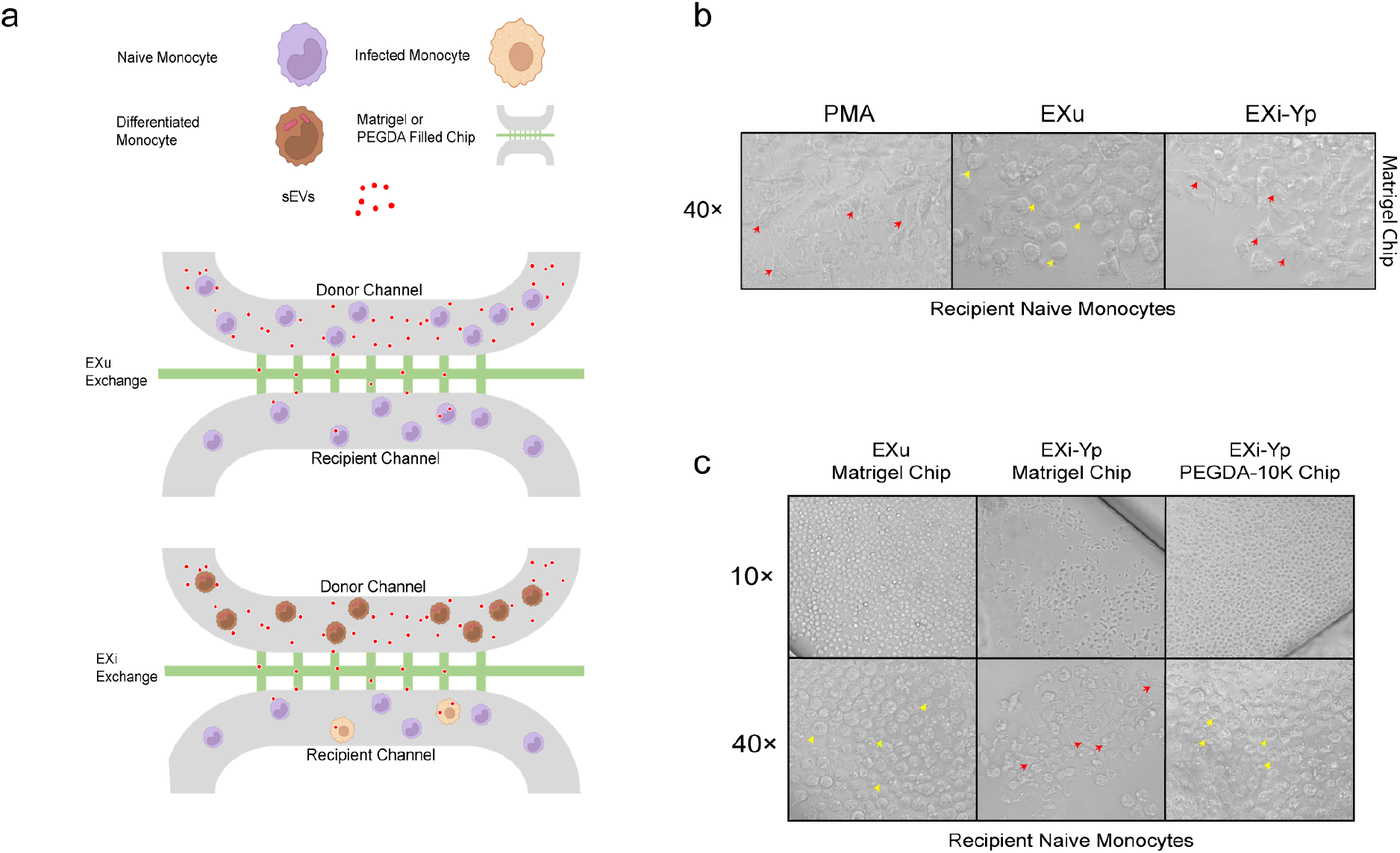
Real-time intercellular exchange of EXi-Yp induces the monocyte to macrophage differentiation phenotype. (a) Illustration of the microfluidic chip experiment to analyze the effect of cell-cell exchange of EXi-Yp on naive recipient monocytes. (b) Differentiation effects of translocated EXi-Yp on naive recipient U937 cells were analyzed on a Matrigel chip. For comparison, a Matrigel chip with an equivalent number of EXu placed in the donor channel, and also another with PMA in the donor channel, were analyzed side by side. Red arrows show examples of differentiated monocytes, while yellow arrows show examples of undifferentiated monocytes. (c) Either uninfected monocytes (EXu panels), or Yp-infected monocytes (EXi-Yp panels), were injected in the donor channel, and the differentiation of naïve monocytes in the recipient channel was analyzed. A control chip with PEGDA-10K as the hydrogel barrier instead of Matrigel was also analyzed side by side. Red arrows show examples of differentiated monocytes, while yellow arrows show examples of undifferentiated monocytes

### EXi-Yp Induce Strong Production of IL-6 In vivo and Confer Protection against Yp Challenge in a Mouse Model of Plague

We used an established mouse model of plague to provide *in vivo* evidence of the IL-6-dependent protective effects observed for EXi-Yp. Groups of mice were administered via the intravenous route (IV) equivalent numbers of either EXu or EXi-Yp, or an equivalent volume of the buffer vehicle (PBS), and their sera were collected both pre-treatment and at 24 hours post treatment for quantitation of proinflammatory cytokines. A panel of mouse pro-inflammatory cytokines very similar to the one used for our *in vitro* studies was used for this analysis. Consistent with the *in vitro* data, treatment of mice with EXi-Yp resulted in a significant increase of IL-6 and IL-10 from this panel (Fig. 11a); IL-8 was not present in the panel because, unlike humans, mice do not carry the IL-8 gene. We next analyzed whether EXi-Yp treatment protects against Yp challenge in mice. Groups of mice were administered through the intravenous route (IV) with either EXi-Yp or with an equivalent volume of the buffer vehicle (PBS) and after 24 hours received a sub-lethal dose of Yp inoculation via the subcutaneous (sc) route (Fig. 11b). Following our IACUC-approved clinical scoring criteria for Yp studies, the two cohorts were monitored daily for 9 days for appearance of disease phenotypes, with mice that started to show strong indications of disease monitored twice a day (Fig. 11b). The results showed that while all the mice in the control group showed strong signs of illness starting on day 8, the EXi-Yp treated mice remained healthy for the duration of the study (Fig. 11b). Together, these *in vivo* findings support the relevance of our *in vitro* results, by demonstrating that EXi-Yp play a protective role by regulating innate immune response to infection with Gram-negative pathogens.

**Figure 11.**
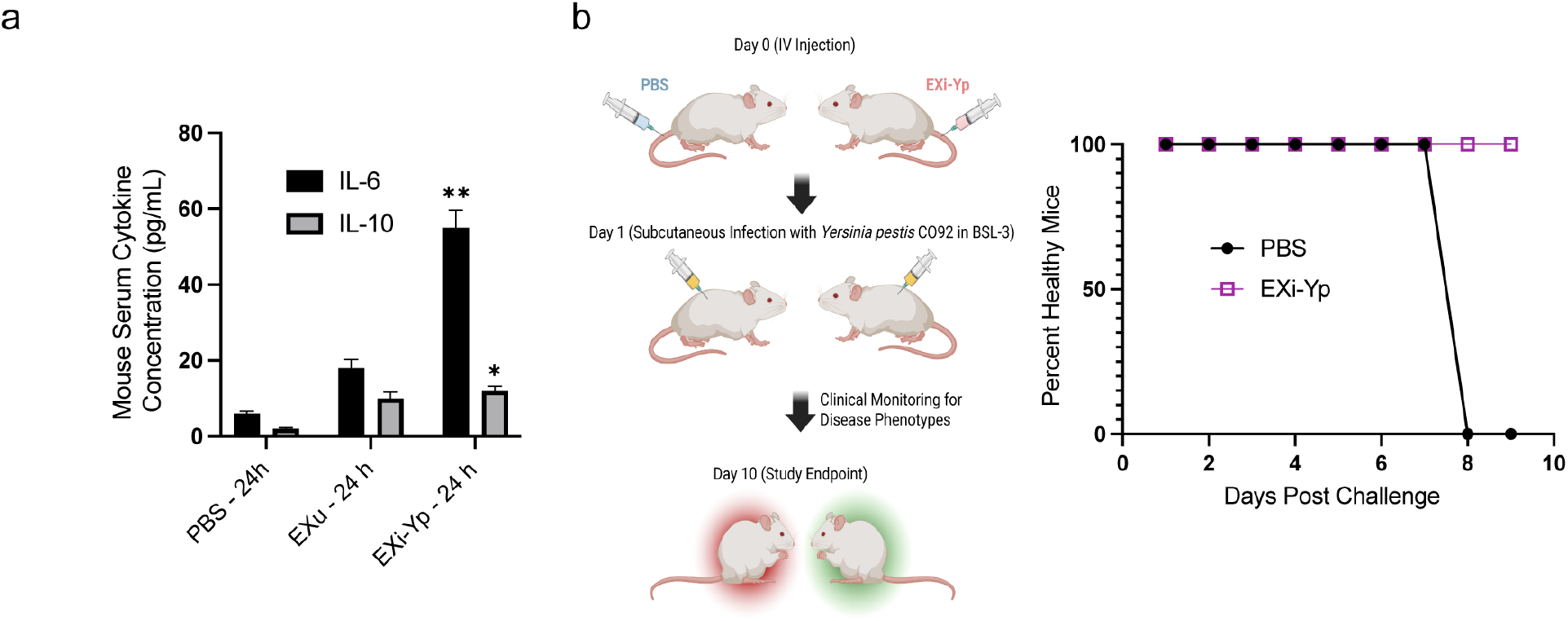
*In vivo* induction of IL-6 by EXi-Yp and its protective effects against Yp challenge in a mouse model of plague. **(a)** BALB/c mice were injected intravenously either with PBS (vehicle control), or with equivalent numbers of EXu or EXi-Yp, and serum samples were collected 24 h post injection for quantitative multiplex probing of pro-inflammatory cytokines. The levels of cytokines significantly induced by EXi-Yp are shown (mean ± SEM, n=5). **(b)** BALB/c mice (n=10 per group) were injected intravenously with either EXi-Yp or with an equivalent volume of the PBS buffer (vehicle control). After 24 hours, the mice were subjected to a sub-lethal dose of Yp CO92 inoculation via the subcutaneous route in our BSL-3 biocontainment facility. For a period of 9 days post challenge, mice were monitored daily (or twice a day after mice showing strong sings of illness) for various clinical phenotypes of disease .

## Discussion

Using infection of human monocytes with Yp and Bt as Gram-negative infection models, we have helped address the continuing significant gap in knowledge of the molecular mechanisms that are engaged by sEVs to regulate innate immunity during infection with Gram-negative bacteria. Quantitative proteomic (RPPA) analysis of cell signaling events at multiple times post EXi-Yp treatment of U937 cells showed strong modulation of a distinct set of host pathways, including activation of p38 MAPK and down-regulation of Jak2 (Table 1). A similar strong activation of p38 MAPK activity was also observed in EXi-Bt-treated monocytes. Related to the EXi-induced effects that we have observed, the activation of p38 MAPK has been shown to induce G1 arrest ^6^, trigger monocyte-to-macrophage differentiation ^52^, cause release of pro-inflammatory cytokines by monocytes ^53–55^, and activate phagocytosis ^56^. Consistent with these established roles for p38 signaling, chemical inhibition of p38 activity dramatically reversed the EXi-associated phenotypes of monocyte to macrophage differentiation and increased IL-6-dependent bacterial clearance. Previous studies have demonstrated that attached populations of monocytes with increased CD68 expression have increased levels of p38 activity, and that chemical inhibition of p38 function reduces the expression of CD68 and adhesion molecules ^57^, corroborating our results that p38 activity regulates EXi-associated monocyte differentiation. We have observed that EXi-induced IL-6 expression is also regulated by increased p38 signaling. These observations may be related to the ability of p38 to activate NF-κB, whose activation has been associated with the EXi-associated phenotypes that we have observed ^57^. As we have shown here that EXi-induced p38 activation in monocytes is a common mechanism of host response to Gram-negative bacteria, it is worth noting that p38 activation has also been reported in monocytes and macrophages treated with sEVs derived from cells infected with *Mycobacterium tuberculosis* or *M. bovis,* ^58,59^, suggesting that EXi-induced p38 activity may be a common signaling regulation of innate immunity across various infections involving either Gram-negative or Gram-positive pathogenic bacteria. Future studies to analyze contributions of other EXi-induced signaling modulations that we have observed should further enhance our understanding of innate immune regulation by sEVs during infection with pathogenic bacteria. For instance, our RPPA results have demonstrated strong deactivation of Jak2, which has been shown to occur during monocyte to macrophage differentiation ^60^ and cause G1 arrest ^61^, suggesting that the Jak2 pathway may be another main target of EXi-Yp regulation. This may relate to the observation that while the EXi-associated monocyte to macrophage differentiation is reversed significantly in the absence of p38 activity, nevertheless some residual level still remains that may involve modulation of Jak2 function.

We have shown that in addition to increased p38 activation and IL-6 release, both EXi-Yp and EXi-Bt also regulate bacterial spread by influencing intracellular bacterial growth/survival. Furthermore, we have found that this increased bacterial clearance depends on p38-dependent induction of IL-6 release. Consistent with these in vitro findings, we have also shown that EXi-YP induce significant increase of IL-6 release *in vivo* and provide protection against Yp challenge in a mouse model of plague. Further support of our *in vitro* findings is provided by the monocyte-to-macrophage differentiation phenotype that is observed following intercellular EXi exchange under conditions that are more physiologically relevant. For these studies, we used our reported microfluidic chip platform that allows functional interrogation of EV effects through real-time cell-cell exchange studies. Interestingly, reduced intracellular bacterial load with sEVs released from cells infected with *M. tuberculosis* has also been demonstrated ^62,63^, and sEV-associated increase in survivability of mice infected with *S. pneumoniae*, another Gram-positive pathogen, has been reported as well ^64^. Therefore, our findings suggest that some of the regulatory pathways engaged by the sEVs to provide enhanced protection may be common between infection with Gram-negative and Gram-positive bacterial pathogens.

Our studies of the EXi effects on naïve monocytes have allowed for construction of a mechanistic model for how they regulates innate immunity through the monocyte-macrophage lineage to protect against infection with Gram-negative pathogens. This model provides a framework for further investigations into not only how EXi influence the progression of infection with bacterial agents but also more broadly what aspects of EXi mechanisms of action may be disease specific. According to our model (Fig. 12), as part of a coordinated response to combat infection, infected monocytes release sEVs that are loaded with specific cargo into the extracellular environment. We propose that the released EXi induce immune responses in both local and distant naïve monocytes, priming them to fight infection more efficiently once they encounter the bacteria. These effects include triggering monocyte differentiation to macrophages that involves increased p38 activity, and p38 induction of IL-6 that in turn leads to increased bacterial clearance to provide a protective effect (Fig. 12). In this regard, we speculate that EXi-induced release of IL-6 and IL-8 (EXi-Yp) may also promote macrophage migration to the site of infection since the production of these cytokines has been shown to recruit nearby immune cells to the site of infection ^65^, an aspect that can be addressed in future studies. Furthermore, future studies can address the potential role of IL-10. Consistent with its demonstrated role ^36^, the increased production of this anti-inflammatory cytokine by EXi-Yp may serve to maintain immune balance, regulating the levels of IL-6 and IL-8 in order to prevent the occurrence of a harmful cytokine storm.

**Figure 12.**
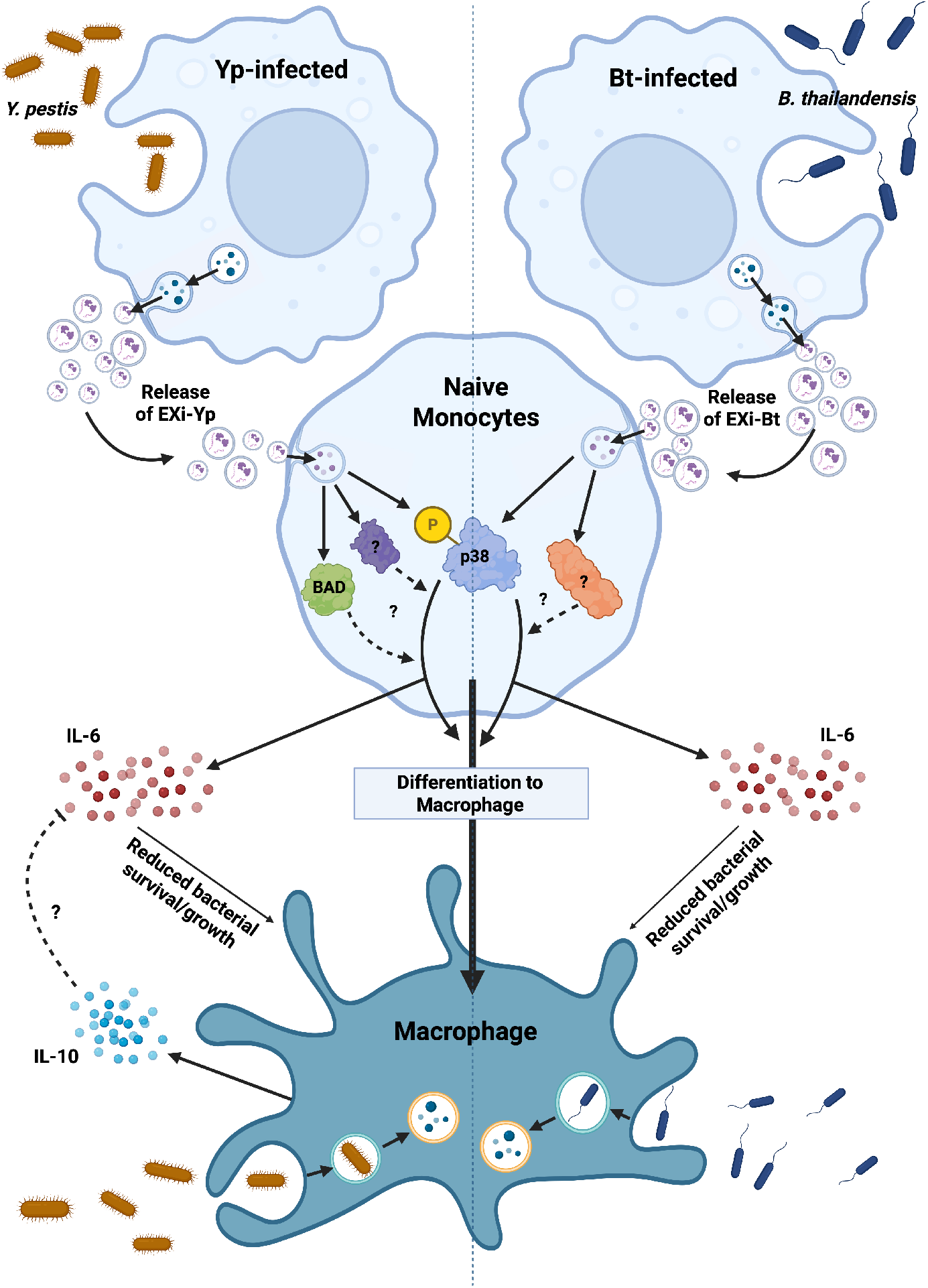
Proposed model of sEV regulation of innate immune response during infection with Gram-negative bacteria through engagement of the monocyte-macrophage lineage. sEVs carrying some specific cargos, including bacterial components, are released from monocytes infected with a Gram-negative bacterial pathogen (such as Yp or Bt), and are internalized by local and distant naïve monocytes. Internalized sEVs modulate specific protein signaling pathways, including the activation of p38 MAPK. Increased p38 activity in naïve recipient cells induces monocyte-to-macrophage differentiation and increased IL-6 release. Elevated IL-6 release in turn allows increased bacterial clearance in the event the cells encounter the bacteria, providing protection against infection. Created in BioRender. Matulis, G. (2025) https://BioRender.com/dfb7ned.

Given the extensive characterizations of Yp and Bt virulence factors and infection mechanisms, and the recent evidence that Yp manipulates EV biogenesis in human neutrophils ^19^, these pathogens provide strong model systems to continue dissecting how bacterial infections shape host immunity through EV-mediated pathways. These future *in vitro and in vivo* studies will further deepen our molecular understanding of EV-based immune regulation, illuminating previously unrecognized aspects of host response to infection that could lay the foundations for developing novel and effective countermeasures.

## Methods

### Bacterial Strains and Cultures

The *Yersinia pestis* (Yp) strain used in this study was CO92 (Pgm-, Pla-), an attenuated version that is pigmentation (pgm)-deficient and carries a frameshift mutation in the pla gene, rendering it non-functional ^66^. This strain carries a fully functional Type Three secretion system (TTSS) ^67^. We also made use of a Yp bacterial strain with the CO92 (Pgm-, Pla-) background lacking the pCD1 plasmid and thus lacking a functional T3SS, generously provided by Dr. James B. Bliska at the Geisel School of Medicine at Dartmouth. For the Burkholderia studies, *B. thailandensis* (Bt) strain E264 was used, which is closely related to *Burkholderia pseudomallei* (Bp) and serves as a well-accepted model for studies of Bp at the BSL-2 level^49^. Bacterial cultures were prepared as described ^66,68^.

### Infection with Yp and Bt

The human monocytic cell line U937, was used in this study. Cells were grown in RPMI 1640 (Lonza) supplemented with 10% Fetal Bovine Serum (VWR) and 2 mM L-glutamine (VWR). EV-depleted Media (EDM) was prepared as described ^17,69^; FBS diluted 1:1 in RPMI was centrifuged at 120,000 × *g* for 3 hours to deplete serum EVs. Culture media of the cells were replaced with EDM 24 hours prior to infection. For infections with Yp, cells were cultured to a concentration of 5 × 10^5^ cells/ml in 100 ml (U937), and incubated with Yp at MOI 10 for 2 hours at 37 °C. After infection, cell cultures were treated with 50 μg/ml Gentamicin (VWR) for 1 hour at 37 °C to eliminate extracellular bacteria and washed three times in PBS (Invitrogen). Cells were then reseeded in 100 ml of EDM supplemented with the maintenance concentration of 5 μg/ml Gentamicin. For Bt infections, cells were incubated with Bt at MOI 1 for 45 minutes at 37 °C. Extracellular bacteria were eliminated by treatment for 1 hour with 75 μg/ml Kanamycin (ThermoFisher). The cells were than washed three times in PBS, followed by reseeding in 100 ml of EDM supplemented with the maintenance concentration of 25 μg/ml Kanamycin

### sEV Purification

sEVs were purified from culture supernatants in a sterile fashion at 48 hours post-infection following the MISEV guidelines ^70^ and as outlined in Fig. S1. Briefly, sEVs were purified by differential centrifugations, which included passage of the supernatant from the high-speed spin (10,000 x g) through a 0.2 μm filter before ultracentrifugation and further purification of the crude sEV pellet by sucrose density gradient centrifugation at 117,000 ×*g* for 3 hours at 4 °C (Fig. S1). A step-wise density gradient was created by layering 2 M, 1.3 M, 1.16 M, 0.8 M, 0.4 M, and 0.2 M sucrose diluted in PBS. The resulting fractions were harvested, diluted in PBS, and pelleted at 117,000 ×*g* for 2 hours at 4 °C. Resulting pellets were resuspended in PBS containing 1× Halt protease inhibitor cocktail (Thermo-Fisher Scientific), filter sterilized using a 0.22 μm syringe filter (Thermo-Fisher), and stored at -80°C. Each fraction was then probed for CD63, TSG101, and Flotillin-1 by western blot. Further exceeding standard practice in the field, to provide additional verification that, as expected, the purified sEVs are free of either bacterial or viral contaminants, media agar plating and plaque assays were performed.

### Western Blot Analysis

Western blot analysis was performed as described ^16^. The following primary antibodies were used: mouse anti-CD63 (EMD Millipore); mouse anti-TSG101 (BD Biosciences); anti-flotillin-1 (Cell Signaling Technology); mouse anti-Yp F1 (Abcam); mouse anti-CD68 (Abcam); anti-p38 (Cell Signaling Technology); phospho-p38 (Cell Signaling Technology), anti-phospho-HSP27 (Cell Signaling Technology), and anti-actin (Abcam). Dilutions were according to the manufacturers’ recommendations. The membranes were incubated with each primary antibody overnight at 4 °C and then washed and incubated for 1 hour at room temperature with goat anti-mouse HRP-conjugated secondary antibody (Cell Signaling) or anti-mouse HRP-conjugated secondary antibody (Cell Signaling Technology). Protein bands were visualized using SuperSignal West-Femto Maximum Sensitivity Substrate (Pierce) and Clarity Western ECL Substrate (BioRad). Images and quantification of the blots were obtained using Chemidoc XRS System (BioRad) and ChemiDoc Gel Imaging System (BioRad). Protein band intensity was quantified using the ImageLab system (BioRad).

### Transmission Electron Microscopy

Ten microliters of each sEV preparation was placed on a copper TEM grid. After 10 sec of absorption, the grid was negatively stained with 2% uranyl acetate. The copper grid was blotted and examined with a FEI Talos F200X TEM, operating at 80 kV.

### Scanning Electron Microscopy

0.1 mg/ml Poly-d-lysine (ThermoFisher Scientific) was added to polycarbonate track-etched (PCTE) membrane filters (Millipore Sigma) and incubated for 1 h at room temperature (RT). After aspiration and drying of the filter, a 40 μl sample volume (1 μl of sEV diluted in 39 μl PBS) was then added and incubated for 10 min at RT. The sample volume was subsequently aspirated and the filter was submerged in a 3.5% glutaraldehyde, 0.5% tannic acid, 0.1 M sodium cacodylate buffer (Electron Microscopy Sciences: EMS) and incubated overnight at 4 °C. The filter was then gently washed with 1 ml of 0.1 M sodium cacodylate buffer, followed by a second wash and then a serial ethanol (EMS) solution exchange of 15%, 30%, 45%, 60%, and then 80% ethanol in 0.1 M sodium cacodylate buffer. Each ethanol dilution was incubated for 10 min at RT and subsequently two 10-minute incubations in 100% ethanol were completed at RT. The filter was then incubated in 50% ethanol: 50% hexamethyldisilazane (HMDS) (EMS) buffer and then 100% HMDS at RT for 10 minutes. The HMDS was aspirated and the filter was left to evaporate at RT. Dried samples were gold sputtered at 20 mA for 20 s using a Denton Desk V and then imaged using a JEOL JSM-7200F set to an accelerating voltage range of 15 kV.

### ZetaView Analysis

Nanoparticle tracking analysis was performed using the ZetaView Z-NTA (Particle Metrix) and its corresponding software (ZetaView 8.04.02). The machine was calibrated according to manufacturer’s protocol. Briefly, 100 nm polystyrene nanostandard particles (Applied Microspheres) were used to calibrate the instrument prior to sample readings at a sensitivity of 65 and a minimum brightness of 20. Automated quality control measurements including, but not limited to, cell quality check and instrument alignment and focus were also performed prior to use. For each measurement, the instrument pre-acquisition parameters were set to a temperature of 23 °C, a sensitivity of 85, a frame rate of 30 frames per second (fps), and a shutter speed of 250. For each sample, 1 mL of the sample diluted in DI water was loaded into the cell, and the instrument measured each sample at 11 different positions throughout the cell, with three cycles of readings at each position. After automated analysis, the mean size (indicated as diameter), the concentration of the sample, and the Zeta potential value, were calculated. Measurement data from the ZetaView were analyzed using the corresponding software, ZetaView 8.04.02, and Microsoft Excel 2016.

### sEV Staining and Treatment with Proteinase K

sEVs were stained using the PKH26 red fluorescent linker kit (Sigma) according to manufacturer’s protocol. Briefly, sEVs were resuspended in the dye solution and incubated for 10 min at 37 °C. The staining was then halted by addition of EDM, and the stained sEVs were pelleted by centrifugation at 117,000 ×*g* for 2 hours at 4 °C. For proteinase K treatment, sEVs were incubated with 500 μg/ml proteinase K (Sigma) for 1 hours at 37 °C, then diluted in PBS and pelleted at 117,000 ×*g* for 2 hours at 4 °C, followed by resuspension in sterile PBS containing 1 × Halt protease inhibitor. The sEV preparations were then filter sterilized using a 0.22 μm syringe filter before further analysis.

### Flow Cytometry Analysis

For flow cytometry analysis of sEV uptake, sEVs were stained as described above using PKH67 fluorescent linker kit (Sigma). U937 cells were incubated with stained sEVs for 6 hours prior to fixation in 2% formaldehyde. Cells were acquired on a FACSCalibur flow cytometer (BD Biosciences) and analyzed using FlowJo 8.8.6 software (Treestar).

### Cell Growth Assays

U937 cells were seeded in 24 well plates at a density of 2.5 × 10^5^ cells/ml in a 1 ml volume of EDM. Yp-infected and Bt-infected cells were prepared as described above. For sEV treatments, cells were incubated with 10 μg of the appropriate sEVs that corresponds to a sEV:Cell ratio of 50:1. At 0-, 24-, 48-, and 72-hours post-treatment, cell concentration and viability were measured by both Trypan blue staining using the Countess Automated Cell Counter (Invitrogen) and by the more sensitive Acridine Orange / Propidium Iodide (AO/PI) staining using the Luna-Fl Dual Fluorescence Counter (VitaScientific). Cell growth rate was calculated as the increase in cell number over each 24-hour period, using the following formula: [Cells_present_ – Cells_past_] / Cells_past_. For cell viability assays, U937 cells were treated as above and at 4-, 6-, 12-, and 24-hours post-treatment cell viability was analyzed using Cell Titer 96 (Promega), according to manufacturer’s protocol. For bacterial growth assays, Yp liquid culture containing 1800 colony forming units (CFUs) were used (calculated by both bacterial plate count and OD_674nm_ measurement). The bacterial cultures were either left untreated or were treated directly with 10 μg of either EXu or EXi, and bacterial CFUs were quantified at 0-, 4-, and 6-hours post-treatment by dilution plating.

### Cell Attachment Assays

U937 cells were seeded at a density of 5 × 10^5^ cells/ml in a 24 well plate and were either left untreated, or were treated with 10 μg of either EXu or EXi that corresponds to a sEV:Cell ratio of 50:1. Untreated cells that were infected with Yp or Bt as described above, were also included. After 24 hours, the culture supernatants were removed and the wells gently washed with PBS. The remaining adherent cells were then trypsinized and quantified by both Trypan blue staining using the Countess Automated Cell Counter and Acridine Orange / Propidium Iodide (AO/PI) staining using the Luna-Fl Dual Fluorescence Counter (VitaScientific).

### Brightfield Microscopy

U937 cells were seeded at a density of 5 × 10^5^ cells/ml in a 12 well plate and were either left untreated, or were treated with 10 μg of EXu or EXi (i.e., sEV:Cell ratio of 50:1), or with 100ng/μl PMA. After 24 hours, the culture supernatants were removed, and wells were gently washed with PBS. Fresh culture media was then added to each well and the cells were incubated for another 72 hours before imaging. Images were obtained using a BioTek Cytation 5 with a 40× objective magnification.

### Yersinia pestis Uptake and Clearance Assays

A total of 5 x 10^5^ U937 cells suspended in 1 mL EDM were added to each well of a 12-well plate and were either left untreated or pre-treated with 10 μg (i.e., sEV:Cell ratio of 50:1) of EXu or EXi-Yp, followed by incubation at 37°C for 24 hours prior to infection with Yp as described above. Uninfected and untreated cells were also included as additional controls. Following treatment with 50 μg/ml Gentamicin for 1h at 37°C, cells were washed twice in EDM and seeded in a 48-well plate in the presence of 5 μg/ml Gentamicin. At 0- and 24-hours post-infection, equivalent numbers of cells per treatment were centrifuged at 800 ×*g* for 5 min at room temperature. The resulting cell pellets were washed in PBS and then lysed in 0.2% Triton X-100 (Sigma-Aldrich), which we have validated to leave bacterial viability intact. The resulting lysates were serially diluted and plated in technical duplicates on Heart Infusion Broth agar plates (ThermoFisher Scientific) with 2% D-xylose and 2.5 mM CaCl_2_. Following incubation at 37 °C to allow colony growth, CFU counts were performed.

### Burkholderia thailandensis Uptake and Clearance Assays

A total of 5 x 10^5^ U937 cells suspended in 1 mL EDM were added to each well of a 12-well plate and were either left untreated or pre-treated with 10 μg (i.e., sEV:Cell ratio of 50:1) of EXu or EXi-Bt, followed by incubation at 37°C for 24 hours. Subsequently, the cells remaining in suspension were discarded and the attached cells were harvested (2x washes with PBS, followed by addition of trypsin and incubation for 5 min at 37°C, 5% CO_2_). The concentrations of the recovered attached cells were determined by the AO/PI stain using the Luna-FL Dual Fluorescence Counter (VitaScientific), and the cells were subsequently infected by Bt as described above. The cells were then washed 2x with PBS, and incubated with 75 μg/mL of Kanamycin in 1 mL PBS for 1 hour, which we have verified to eliminate the extracellular bacteria. The cells were then washed 2x in PBS, and half of the cells were immediately lysed by incubating in 1 mL of 0.2% Triton-X100 (Sigma-Aldrich) for 5 minutes at 37°C. The resulting cell lysates were serially diluted and each dilution was plated in technical duplicates on LB plates and following incubation at 37 °C to allow colony growth CFU counts were performed. The remaining half of the cells from each treatment were resuspended in EDM containing the maintenance concentration of 25 μg/mL kanamycin and returned to the 12-well plate to incubate for 24 hours. During this incubation period, the wells were spiked twice with Kanamycin at 8- and 16-hour time points to remove any remaining extracellular bacteria without harming the U937 cells. Subsequently, the cells were harvested for CFU counts as described above.

### Multiplex Cytokine Arrays

In a 24-well plate, U937s were seeded at a density of 5 × 10^5^ cells/ml in a 2 ml volume of EDM per treatment. To each treatment well, 10 μg of either EXu or EXi (i.e., sEV:Cell ratio of 50:1) was added at 0 h. Untreated cells that were infected with Yp or Bt as described above (positive control), and untreated and uninfected cells (baseline control), were also included. At 12, 24, or 48 h post treatment, 500 μl of the cell culture supernatant was harvested from each well and cleared by centrifugation at 800 ×*g* for 5 min at room temperature. The resulting supernatants were then used for cytokine measurements using the Aushon Cirscan System (Aushon Biosystems), following the manufacturer’s instructions. First, the 10-plex human inflammatory cytokine array from Aushon Biosystems was analyzed in biological duplicates. This panel allows multiplex quantitation of the human pro-inflammatory cytokines IFNγ, IL-1α, IL-1β, IL-2, IL-4, IL-6, IL-8, IL-10, IL-12p70, and TNFα. Next, based on the results of the 10-plex analysis for EXi-Yp, three biological repeats of the cytokine measurements were performed using a 3-plex Aushon Ciraplex Human Cytokine Array for IL-6, IL-8, and IL-10, following the manufacturer’s protocol.

### IL-6 ELISA Measurements

Untreated and treated U937 cells were transferred to microcentrifuge tubes and pelleted at 800 x g for 5 minutes. Each resulting supernatant was removed and transferred to a new microcentrifuge tube. 1X Halt Protease Inhibitor cocktail (ThermoFisher Scientific) was added and the samples were stored at -80°C until performance of ELISA measurements using the Legend Max Human IL-6 ELISA Kit (BioLegend 430507) according to manufacturer’s recommended procedures.

### siRNA Studies

U937 cells were seeded at a density of 2.5 × 10^5^ cells/ml in a 1 ml volume of EDM per treatment. For qRT-PCR validation of knockdowns, cells were treated with five nMol of pre-validated siIL-6 or siIL-8 siRNAs (Thermo-Fischer) in complex with Lipofectamine 2000 (Thermo-Fischer) following manufacturer’s protocol, and incubated at 37 °C for either 24 h or 48 h. As control, cells treated with the validated negative control siRNA no. 1 (Thermo-Fischer) were also included side by side. Total RNA was then extracted using the RNeasy Mini kit (Qiagen) and cDNA was generated using the All-in-One cDNA Synthesis SuperMix (Biotool). The qRT-PCR reaction was prepared using SYBR Select Master Mix (Thermo-Fischer) and pre-validated primers for GAPDH, or IL-6, or IL-8 (Realtimeprimers), following the manufacturer’s protocol. Data was analyzed using the ΔΔCt method ^71^. To analyze recipient cell response to EXi after either IL-6 or IL-8 knockdown, validated knockdown conditions were used to treat U937 cells with either siIL-6, siIL-8, or control scrambled siRNA no. 1 (5 nMol siRNA; 24 h treatment), and the cells were subsequently left either untreated (control) or were treated with equivalent numbers of EXu or EXi-Yp for 24 h. Effects on rate of growth, cell attachment, and bacterial clearance were analyzed.

### p38 Chemical Inhibitor Treatment

U937 cells were cultured in EDM RPMI containing 500 nM of the p38 inhibitor PH797804 (APExBIO) and were allowed to incubate for 1 hour at 37°C and 5% CO2. The cells were then either left untreated or were treated with equivalent numbers of EXi-YP or EXi-Bt, followed by further manipulations described above for measurements of cell growth rate, cell attachment, and IL-6 levels. Control wells that contained an equivalent volume of DMSO, the solvent in which the p38 inhibitor was reconstituted, were run in parallel.

### Mass Spectrometry Analysis of EXi-Yp

sEV lysate samples were prepared and MS analysis was performed as previously described ^16^. Tandem mass spectra collected by Xcalibur (version 2.0.2) were searched against the NCBI *Yersinia pestis* protein database using SEQUEST (Bioworks software from ThermoFisher, version 3.3.1) with full tryptic cleavage constraints, static cysteine alkylation by iodoacetamide, and variable methionine oxidation. Mass tolerance for precursor ions was 10 ppm and mass tolerance for fragment ions was 0.5 Da. The SEQUEST search results were filtered by the criteria “Xcorr versus charge 1.9, 2.2, 3.0 for 1+, 2+, 3+ ions; ΔCn > 0.1; probability of randomized identification of peptide < 0.01”. Confident peptide identifications were determined using these stringent filter criteria for database match scoring followed by manual evaluation of the results.

### RPPA Proteomic Profiling

The quantitative RPPA analysis of phosphorylation and total protein level changes in response to sEV treatment was performed as described ^27–29,72^. Sample slides were probed with 173 different antibodies against either total or phosphorylated forms of proteins that are involved in various cell signaling pathways, including those that are known to be relevant to infectious diseases. A complete list of all the antibodies used for this analysis, categorized by functional relevance, is provided in Supplementary Table 1. For each specific post-infection or post-treatment time point, the RPMA results for the different treatment conditions were normalized to the control group (uninfected and untreated) in order to calculate relative fold-change levels. For total protein levels, a 2-fold increase/decrease was set to be significant and for phosphorylation levels the significant fold increase/decrease was set to 1.5. Extensive literature searches were performed using Pubmed for all the proteins that showed significant change to assess the significance of the expression fold change.

### Y. pestis lipid A purification

Lipid A from Y. pestis was purified using a modified version of a previous method ^73^. Briefly, lyophilized whole bacterial cell lysate, and EVs from infected or noninfected host cells, prepared as described above, were re-suspended in 3.8 mL CHCl3:MeOH:water (1:2:0.8, v/v/v) in 15-mL PTFE-lined cap glass vials (Supelco, Sigma-Aldrich, St. Luis, MO). Following agitation for 1 h by rocking at room temperature (RT), centrifugation at 1000 x g for 10 min at RT, supernatants were transferred to fresh 15-mL PTFE-lined glass vials, and pellets were re-extracted with the same solvent mixture. After organic extraction, vials were centrifuged and supernatants were combined. The resulting pellets were dried under N2 gas stream to remove any remaining solvent. The pellets were re-suspended in 10 mL 50 mM sodium acetate, pH 4.5, and vortexed for 5 min. The samples were placed on a heating block at 100°C and incubated for 2 h. Afterwards, the samples were cooled to RT, and 4 mL CHCl3:MeOH (2:2, v/v) was added and vortexed for 5 min. The samples were partitioned by centrifugation at 1000 x g for 20 min at RT. The lower phase was transferred to a fresh 7-mL PTFE-lined cap glass vial, and 3 mL CHCl3:MeOH (2:1, v/v) was added to the upper phase for re-extraction. This step was repeated twice. The three resulting lower phases were combined and dried under N2 gas. Finally, 5 mL hexane was added to the aqueous phase, centrifuged 1000 x g for 20 minutes, the upper (hexane) phase was pooled together with the dried, combined lower phases, and stored at -80°C. Samples were first dissolved in 100 µL 100% CHCl3 by 1 min vortex and 1 min sonication. Subsequently, 200 µL 100% MeOH was added for a final ratio of CHCl3:MeOH (1:2, v/v) prior to LC-MS/MS analysis.

### Liquid Chromatography-Tandem Mass Spectrometry (LC-MS/MS) Analysis for Detection of Yp Lipid A

Liquid chromatography (LC) was performed on a Dionex Ultimate 3000 UHPLC (Thermo Fisher Scientific) with a column temperature of 45°C and a flow rate of 0.5 mL/min throughout the entirety of the run. A sample volume of 30 µL was injected onto the Kinetex C18 Evo (2.6µm, 100Å, 150 x 2.1mm) reverse-phase column. The column was pre-equilibrated with 30% solvent B (90% isopropanol, 10% acetonitrile, 0.1% formic acid, 10 mM ammonium formate) and 70% solvent A (60% acetonitrile, 40% water, 0.1% formic acid, 10 mM ammonium formate) before injection. After injection, the column was maintained for 3 min, then increased to 43% solvent B over 3 min, increased to 45% over 0.2 min, 65% over 9.8 min, 85% over 6 min, 100% over 2 min. A plateau of 100% solvent B was maintained for 5 min before a sharp decrease to 30% over 0.1 min. The starting conditions of 30% solvent B was maintained for 2.9 min before the blank injection, which was performed after each sample to clean the column from any residual lipid A species. The blank injection method started at 30% solvent B and was maintained for 1 min before beginning a ramp gradient to 100% solvent B over 5 min. Solvent B remained consistent for 2 min and was then decreased to 30% solvent B over 0.2 min. The equilibration conditions were held for 2.8 min before injection of the next sample. Lipid A analysis was performed on a Q Exactive Plus Hybrid Quadrupole-Orbitrap Mass Spectrometer (Thermo Fisher Scientific) equipped with a Heated Electrospray Ionization (HESI-II) source. Spectra were acquired in negative-ion mode by top-4 data dependent (dd)-MS2 throughout the 32-min run. Targeted-selective ion-monitoring (SIM)/dd-MS2 was from 10 to 30 min. Full MS parameters: resolution 140,000, AGC target 1e6, Max IT 50 ms, and scan range 700 to 1800 m/z. dd-MS2 Parameters: resolution 17,500, AGC target 1e5, MAX IT 75 ms, and (N)CE: 35. Targeted-SIM: resolution 70,000, AGC target 1e5, Max IT 100 ms, and isolation window 4.0. Inclusion list ions m/z: 1097.6866, 1165.6752, 1279.8536, 1307.8856, 1323.8806, 1323.8837, 1454.9430, 1454.9442, 1506.0471, 1534.0792, and 1534.0817. Ionization Parameters: spray voltage 3200, capillary temperature 300°C, S-Lens RF level 60, aux gas heater temp 325°C, sheath gas 40, and aux gas 10.

### Microfluidic Chip Studies

The next-generation microfluidic chips used for our studies were fabricated as described^51^, with the exception of the PEGDA hydrogel formulation. For PEGDA-based chips, the central hydrogel region was prepared using 20% (w/v) PEGDA with an average molecular weight of 10,000 Da, supplemented with 0.5% (w/v) lithium phenyl-2,4,6-trimethylbenzoylphosphinate (LAP) as a photoinitiator, dissolved in water. This formulation replaced the PEGDA 400 and Irgacure photoinitiator used in the original design. For analyzing the effect of EXi-Yp transmigration on the chip to interact with naïve recipient monocytes, Yp-infected and uninfected U937 monocytes were prepared at a concentration of 2 X 10^6 cells/mL and used as described above for purification of sEVs. The donor channel of a chip was loaded with the same numbers of either EXu or EXi-Yp and the recipient channel with naïve recipient U937 monocytes, and 48 hours later the monocyte to macrophage differentiation phenotype was analyzed. Another chip with PMA injected into the donor channel and naïve recipient monocytes in the recipient channel was also analyzed side by side. For the cell-cell exchange studies, uninfected and Yp-infected U937 monocytes maintained in RPMI medium supplemented with maintenance concentrations of gentamycin were seeded into the donor channel, and naïve U937 monocytes were seeded in the recipient channel of a microfluidic chip. The two cell channels were separated by a central hydrogel region of either Matrigel or polyethylene glycol diacrylate (PEGDA)-10, forming a diffusion barrier between donor and recipient compartments. Following cell seeding, chips were incubated under standard cell culture conditions (37 °C, 5% CO₂) for 48 hours. At the end of the incubation period, chips were imaged for analysis.

### Mouse Studies

For cytokine measurements, BALB/c mice (Charles River Laboratories) were injected intravenously with either 50 µg of EXi-Yp or EXu per animal, or with an equivalent volume of the vehicle control (PBS). Five mice were analyzed per group. Serum samples were collected pre-injection, and at 24 hours post injection. They were analyzed using an Aushon Cirascan system (Aushon Biosystems) as described above, using the Aushon Ciraplex Mouse 8-Plex Array that allows multiplex quantitation of the mouse pro-inflammatory cytokines IFNγ, IL-1β, IL-6, IL-10, IL-12p70, IL-17, and TNFα. For the infection challenge study, BALB/c mice were injected intravenously with PBS or 50 µg of purified EXi-Yp (10 mice/group). Subsequently, at 24 hours post injection, mice were infected with a virulent strain of Yp (CO92) in our BSL-3 biocontainment facility via the subcutaneous route. For 9 days post-challenge, the mice were monitored daily (or twice a day for animals that started showing strong signs of illness) using our IACUC-approved clinical scoring protocol for Yp infection. Clinical scoring for appearance, mobility, attitude, response to stimuli, respiratory distress, and body condition were performed.

### Statistical Analysis

Statistical analyses were performed using R studio version 0.99.465 or GraphPad Prism 10.6.1. Normality and homogeneity of variance were tested for all data sets using Shapiro-Wilk and Barlett’s tests respectively. For parametric data, significance was tested using one-way ANOVA with Dunnet’s post-test, and for non-parametric data significance was tested using Kruskal-Wallis with Dunn’s post-test unless noted otherwise.

## Acknowledgements

This work was supported in part by the NIH (NIAID) awards 1R41AI122666-01A1 and 1R42AI122666-03 to RMH, and also by the USMRMC (DoD) Grant W81WH-15-T-0003 to RMH, and the Dean’s Excellence in Research Award to RMH at George Mason University. The funders had no role in the design of the study and collection, analysis, and interpretation of data, or in writing the manuscript. We would like to acknowledge the invaluable assistance of Dr. Rekha Panchal and her group at USAMRIID with collection of mouse sera for cytokine analysis, and the assistance of Dr. Carolina Salvador Morales for the TEM studies of EXi-Bt . We would also like to thank Dr. Remi Veneziano at George Mason University for providing the equipment and facilities necessary to manufacture the microfluidic chips. We are grateful to Kathryn Gerrish, a former student in the Hakami laboratory, for providing assistance with the p38 assays.

