## Supplementary Figures for "Small Extracellular Vesicles from Gram-Negative Bacterial Infections Induce Myeloid Differentiation via p38 Signaling and Coordinate Antibacterial Defense Through the p38–IL-6 Axis"

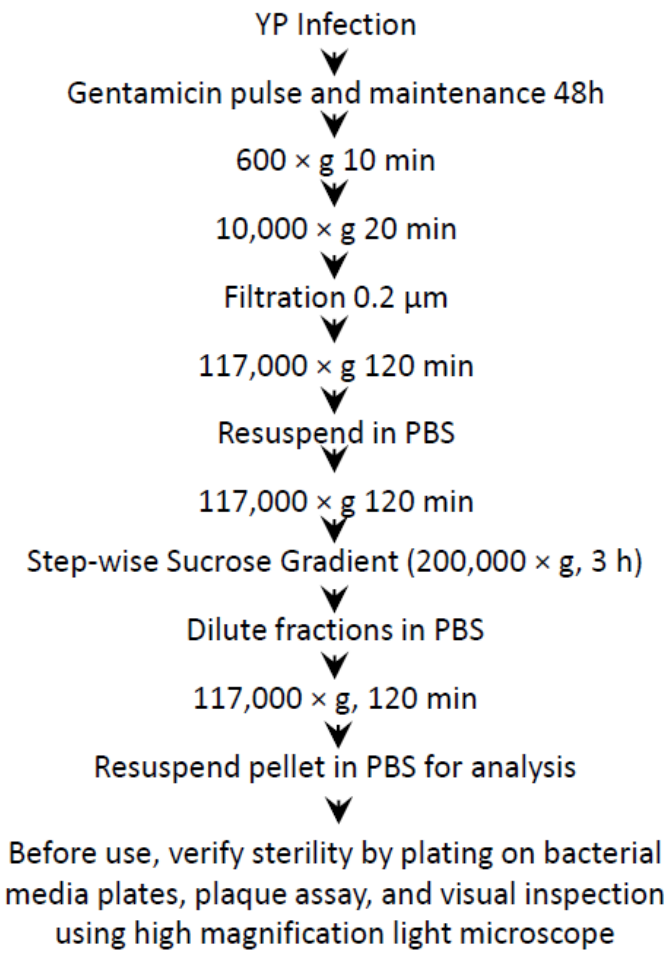


**Figure S1**. Flow chart of sEV purification. For each purification, sEVs from uninfected cells were also recovered and purified side by side using the same procedure.

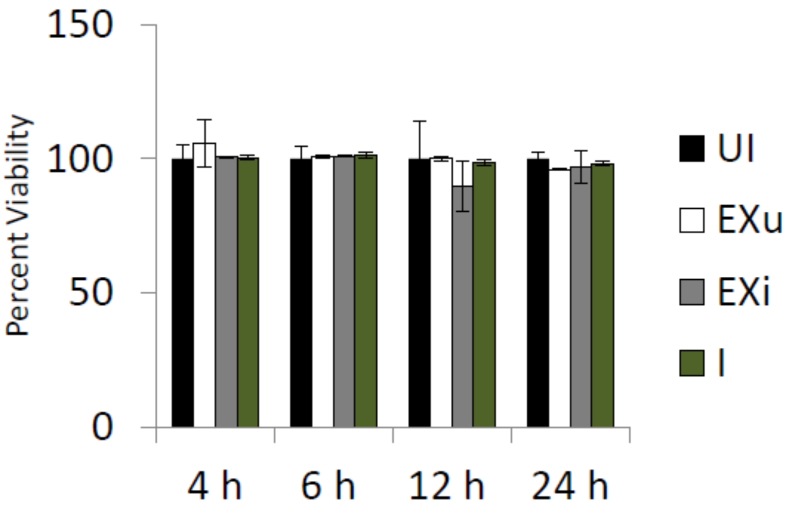


**Figure S2.** EXi-Yp do not affect naïve recipient cell viability. U937 cells were either left untreated, or were treated with equivalent numbers of EXu or EXi-Yp, or were infected with Yp, and cell viability was measured at 4 h, 6 h, 12 h, and 24 h post-treatment using the CellTiter-Glo luminescence cell viability assay (means ± SEM; n=2).

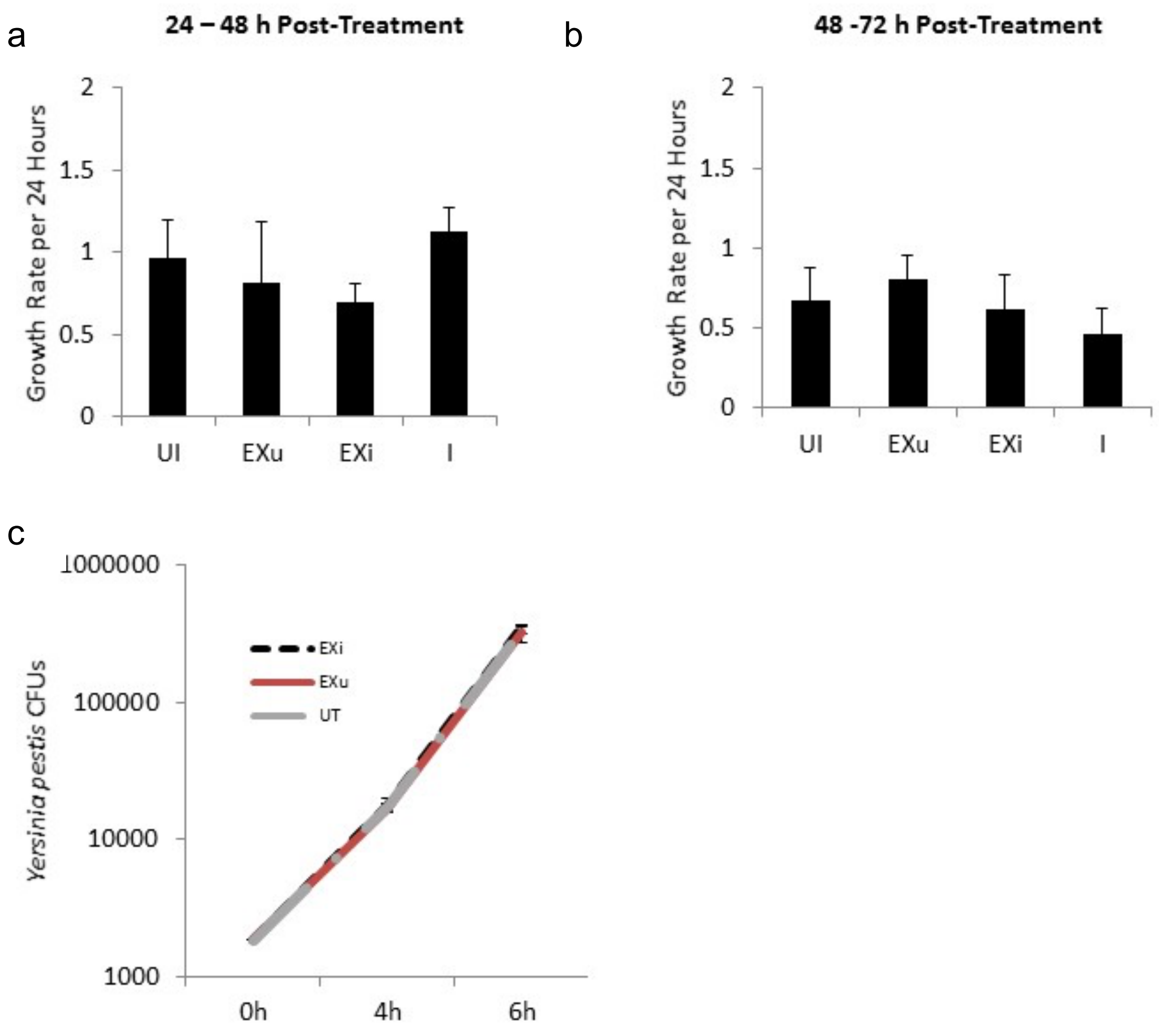


**Figure S3.** Growth rate decrease induced either by EXi-Yp treatment or infection with *Yersinia pestis* does not extend beyond the first 24 h and is a consequence of host response mechanisms. Naïve U937 cells were either left untreated, or were treated with equivalent numbers of EXu or EXi-Yp, or were infected with Yp, and cell growth rate was quantified from **(a)** 24 to 48 h post-treatment and **(b)** 48 to 72 h post-treatment (mean ± SEM; n=5). **(c)** Yp bacteria were either left untreated (grey) or were directly incubated with equivalent amounts of EXu (red) or EXi-Yp (black). Total CFUs were quantified at 0 h, 4 h, and 6 h post treatment by dilution plating of bacteria.

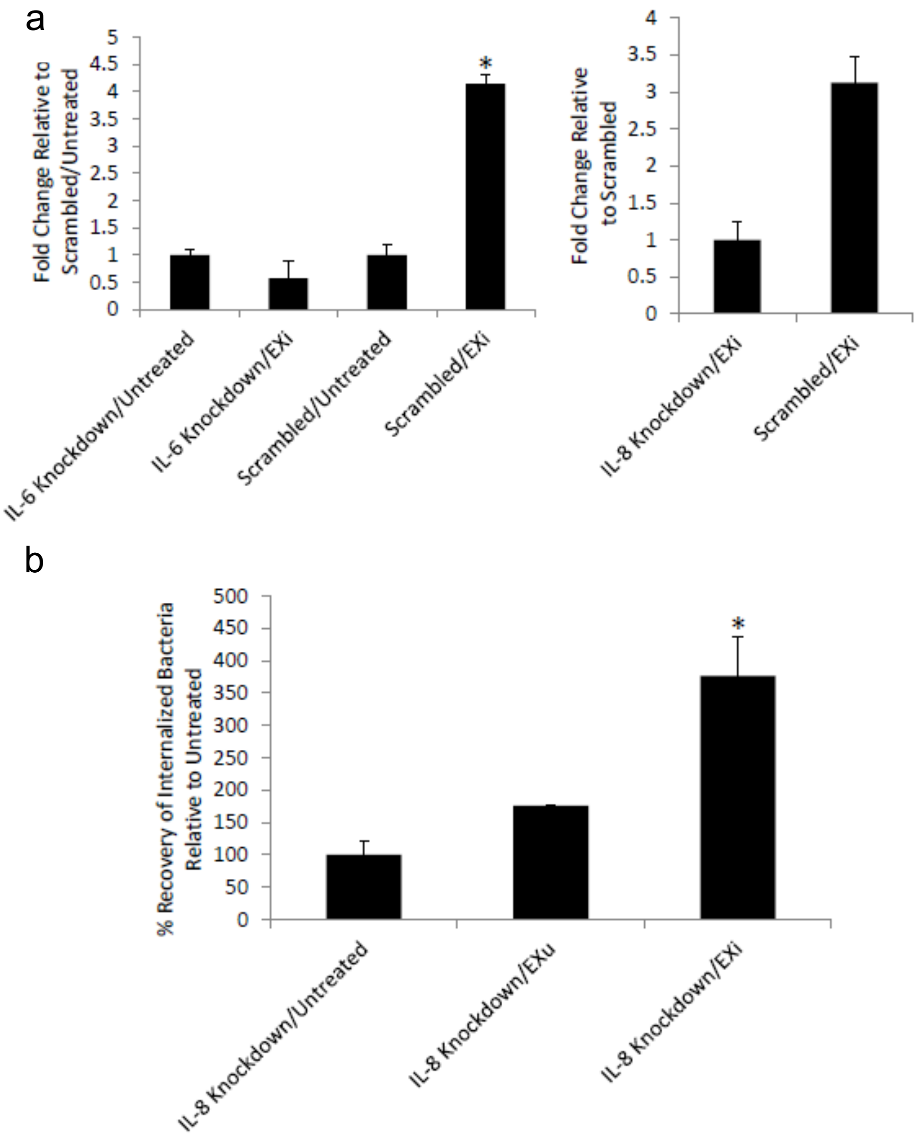


**Figure S4.** Transfection with either siIL-6 or siIL-8 efficiently knocks down IL-6 and IL-8 expression respectively in recipient U937 monocytes, and IL-8 knockdown in recipient cells does not abrogate EXi-Yp induced increase in bacterial uptake. **(a)** Naïve U937 monocytes were transfected with either siIL-6 siRNA, or siIL-8 siRNA, or control scrambled siRNA for 24 h and then either were left untreated or were stimulated with EXi-Yp for 24 h. Intracellular levels of IL-6 or IL-8 mRNA were subsequently quantified by qRT-PCR. Basal IL-8 levels were undetectable by qRT-PCR. (mean ± SEM; n=2). **(b)** IL-8 knockdown does not abrogate increased bacterial uptake by EXi treated cells (mean ± SEM; n = 2).

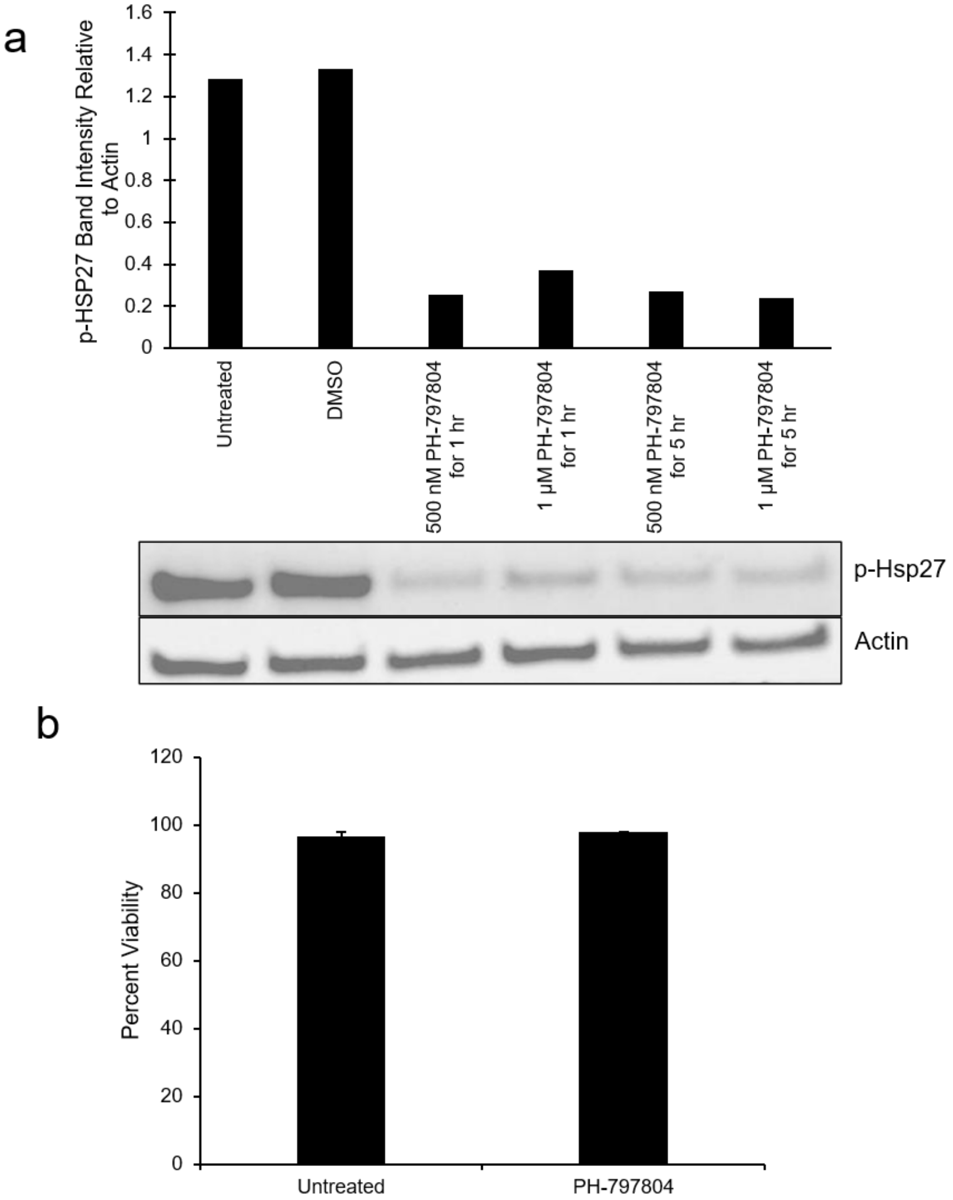


**Figure S5. (a)** U937 cells were either left untreated, or were treated with DMSO, or with the p38 inhibitor PH-797804 at varying concentrations and for various amounts of time. The cells were lysed and probed for p-Hsp27 expression. Phospho-Hsp27 band intensity was quantified relative to actin. **(b)** U937 cells were either left untreated or treated with the p38 inhibitor PH-797804 and allowed to incubate for 24 h. At 24 h, the viability of the cells was assessed using AO/PI (mean ± SEM, n=3).

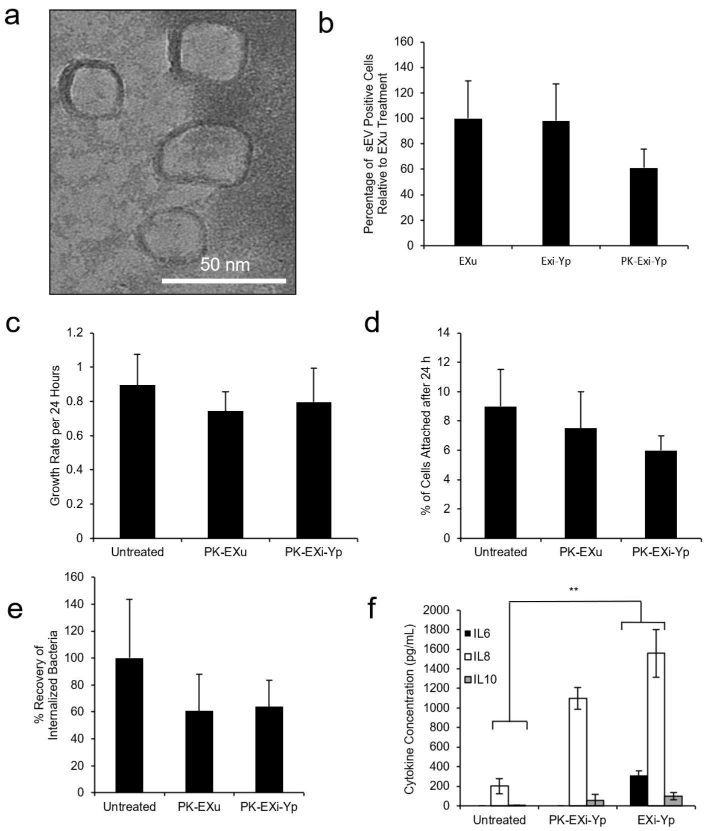


**Figure S6.** sEV surface proteins are important for EXi-Yp induction of monocyte differentiation and IL-6 release. **(a)** TEM of EXi-Yp following treatment with proteinase K (PK-EXi) shows they remain as intact vesicles. **(b)** Comparative flow cytometry analysis of naïve monocytes treated with equivalent numbers of PKH67-stained EXu, EXi, PK-EXu, or PK-EXi shows an overall lower level of uptake for PK-treated vesicles although the difference between PK-treated and untreated vesicles is not statistically significant. The results represent percentage of positive cells relative to EXu (mean ± SEM n=4) **(c)** Naïve U937 cells were either left untreated or were treated with equivalent numbers of PK-EXi or PK-EXu, and cell growth rate was quantified from 0 to 24 h post treatment (mean ± SEM, n=3). **(d)** Naïve monocytes were either left untreated or were treated with equivalent numbers of PK-EXi or PK-EXu for 24 h. Following removal of non-adherent cells, the number of adhered cells was quantified using trypan blue staining assay (mean ± SEM, n=2), **(e)** Naïve U937 cells were either left untreated or were pre-treated with equivalent numbers of PK-EXi or PK-EXu for 24 h, and subsequently infected with Yp. Internalized bacteria were quantified by CFU count at 0 h post infection and percent recoveries of internalized bacteria relative to untreated uptake were measured (mean ± SEM, n=3). **(f)** Naïve human monocytes were either left untreated or treated with equivalent numbers of EXi or PK-EXi. The levels of IL-6, IL-8, and IL-10 in culture supernatants were then quantified at 48 h post treatment using the Aushon Cirascan system (mean ± SEM, n=3).

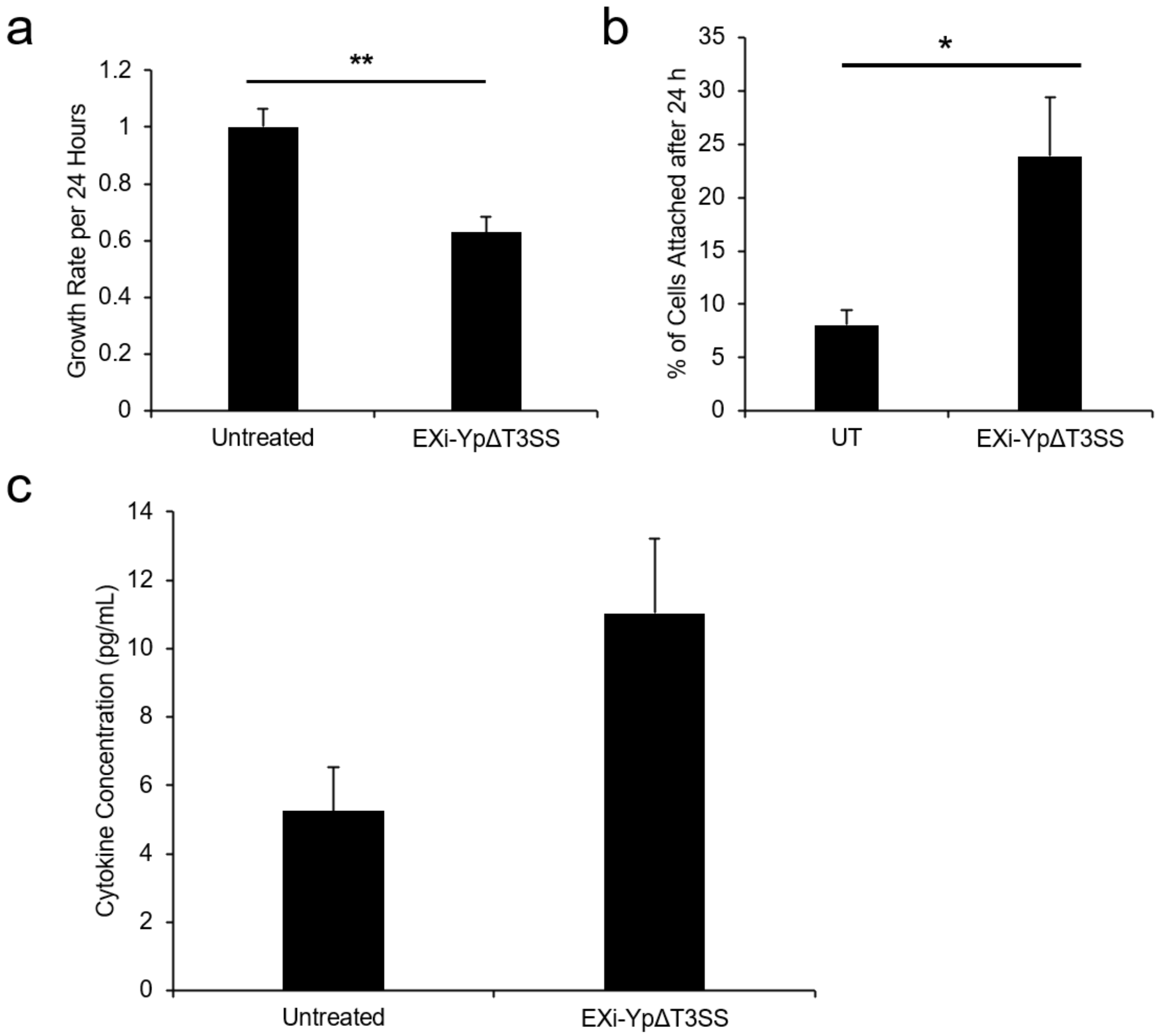


**Figure S7. (a)** Naïve U937 cells were either left untreated or were treated with EXi-YpΔT3SS and cell growth rate was quantified from 0 to 24 h post-treatment (mean ± SEM, n=7). **(b)** Naïve U937 cells were either left untreated or were treated with EXi-YpΔT33SS and after 24 h the number of cells that had adhered were quantified by AO/PI (mean ± SEM, n=3). **(c)** U937 cells were either left untreated or were treated with EXi-YpΔT3SS. After 48 hours, the supernatant was analyzed for IL-6 production (mean ± SEM, n=2).

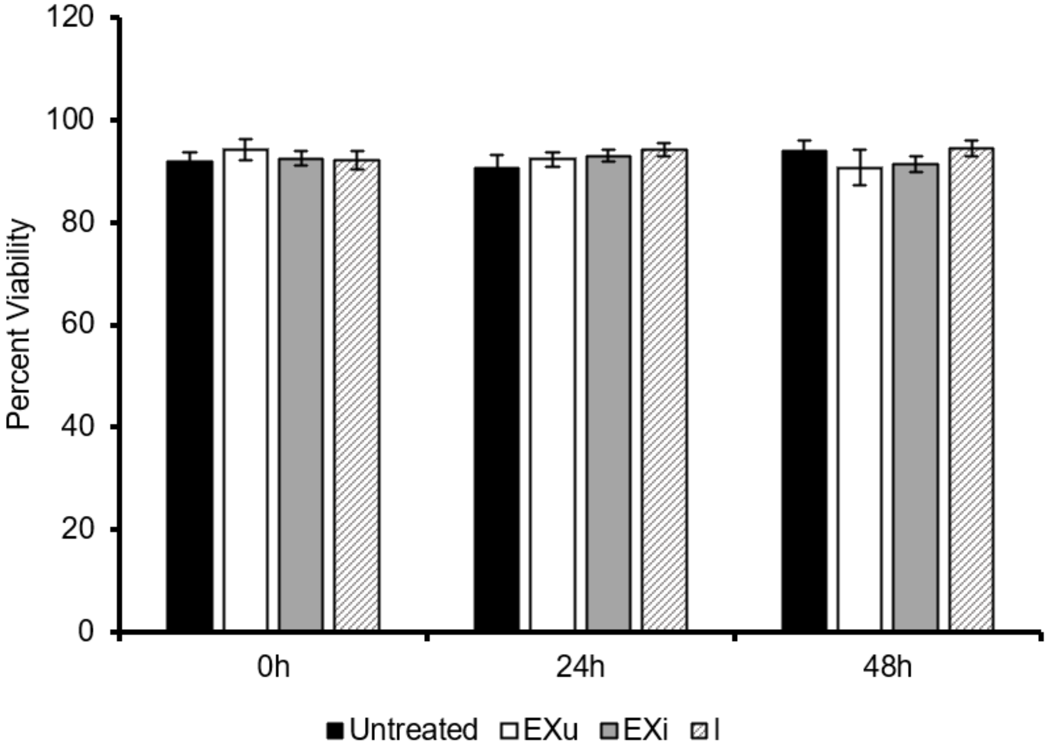


**Figure S8.** Naïve U937 cells were either left untreated, or were treated with equivalent numbers of EXu or EXi-Bt, or were infected with Bt. Cell viability was assayed by AO/PI staining at 0 h, 24 h, and 48 h (mean ± SEM, n=4).

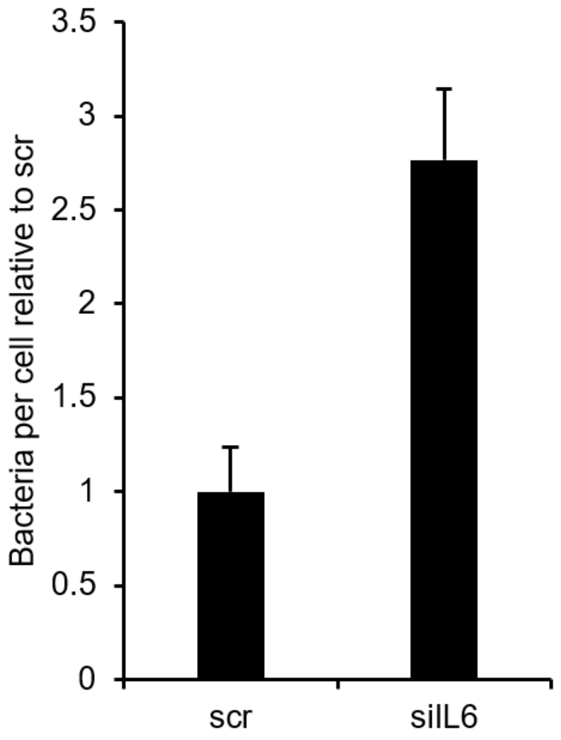


**Figure S9.** U937 cells were treated with either scrambled siRNA or siRNA targeting IL-6 and incubated for 24 hours. U937 cells were then left untreated for an additional 24 hours. The attached cell population was subsequently infected with Bt. The infected cells were lysed 24 hours later, and plated for enumeration of intracellular bacteria (mean ± SEM, n=2)

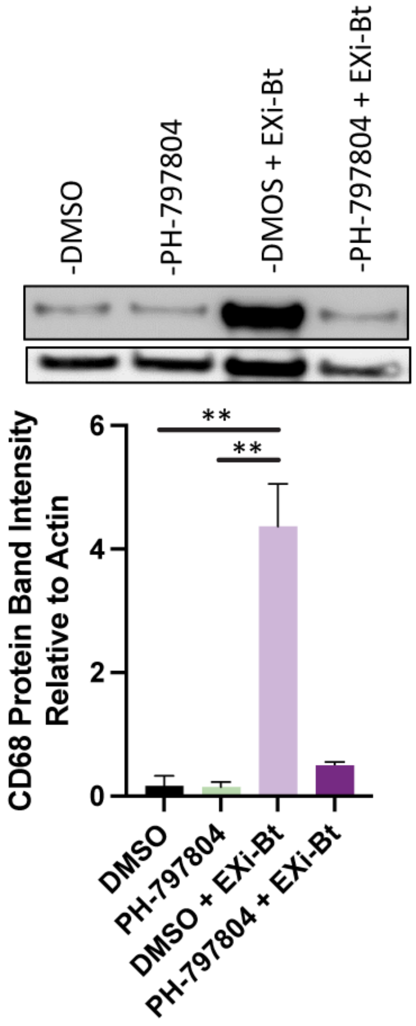


**Figure S10.** U937 cells were pre-treated with either the p38 inhibitor PH-797804 or DMSO, and then were either left untreated or treated with EXi-Bt. After 24 hours, the attached cells were lysed and probed for CD68 expression. The CD68 band intensity relative to actin is shown (mean ± SEM, n=3).

**Table S1.** List of antibodies used for RPPA analysis and their target pathways.

| Protein Name | Pathways Growth/Cell Cycle/Differentiation | Company | Catalog # | Dilutions |
|---|---|---|---|---|
| A-Raf S299 | MAPK; MEK/ERK | CellSig | 4431 | 1:50 |
| ALK | MAPK and PI3 kinase | CellSig | 3633 | 1:50 |
| ALK Y1586 | MAPK and PI3 kinase | CellSig | 3343 | 1:1000 |
| ALK Y1604 | MAPK and PI3 kinase | CellSig | 3341 | 1:50 |
| Androgen Rec S650 | ZIPK/DAPK3 | Millipore | 07-1375 | 1:1000 |
| Androgen Rec S81 | ZIPK/DAPK3 | Abcam | ab47563 | 1:1000 |
| ATM S1981 | DNA damage | CellSig | 5883 | 1:50 |
| ATR S428 | DNA damage | CellSig | 2853 | 1:50 |
| Aurora A/B/C T288/232/198 | Cell cycle/ histone H3 phosphrylation | CellSig | 2914 | 1:500 |
| B-Raf S445 | AKT/PKA | CellSig | 2696 | 1:50 |
| c-Abl T735 * | p53 | CellSig | 2864 | 1:50 |
| c-Met * | Upstream RAS/PI3K/ STAT/Mnt/Notch | Abcam | ab51067 | 1:200 |
| c-Myc * | p38 MAPK/ERK | CellSig | 9402 | 1:100 |
| c-Raf S338 | ERK1/2; Ras | CellSig | 9427 | 1:200 |
| c-Src | AKT/PI3 kinase/p38 MAPK/ MEK;ERK | SantaCruz | sc-18 | 1:200 |
| Chk-1 S345 | DNA damage | CellSig | 2341 | 1:1000 |
| Chk-2 S33/35 | DNA damage | CellSig | 2665 | 1:1000 |
| CREB S133 | cAMP/p38 MAPK | CellSig | 9191 | 1:1000 |
| Cyclin A | Cell cycle regulators | CellSig | 4656 | 1:2000 |
| Cyclin B1 | Cell cycle regulators | CellSig | 4135 | 1:500 |
| Cyclin D1 | Cell cycle regulators | BD | 554180 | 1:1000 |
| EGFR | MAPK/AKT/JNK | CellSig | 2232 | 1:1000 |
| EGFR Y1045 | MAPK/AKT/JNK | CellSig | 2237 | 1:1000 |
| EGFR Y1068 | MAPK/AKT/JNK | CellSig | 2234 | 1:1000 |
| EGFR Y1148 | MAPK/AKT/JNK | CellSig | 4404 | 1:500 |
| EGFR Y1173 | MAPK/AKT/JNK | BioSource | 44-794 | 1:1000 |
| eNOS S113 * | AKT/PKA/AMPK | CellSig | 9575 | 1:1000 |
| eNOS/NOSIII S116 * | AKT/PKA/AMPK | Upstate | 07-357 | 1:1000 |
| Ephrin A3/A4/A5 Y799/799/833 * | Ephrin/Rac | Abcam | 124881 | 1:100 |
| ErbB2 Y1248 * | PI3 kinase/AKT/PLC/ PKC/MAPK/STAT | CellSig | 2247 | 1:1000 |
| ErbB2 Y877 * | PI3 kinase/AKT/PLC/ PKC/MAPK/STAT | Imgenex | IMG-90189 | 1:500 |
| ErbB3 * | PI3 kinase/AKT | CellSig | 4754 | 1:1000 |
| ErbB3 Y1289 * | PI3 kinase/AKT | CellSig | 4791 | 1:1000 |
| ERK 1/2 T202/Y204 * | MAPK | CellSig | 9101 | 1:1000 |
| Etk Y40 | ? | CellSig | 3211 | 1:1000 |
| Ezrin/Radixin/Moesin T567/564/558 * | Rho | CellSig | 3141 | 1:1000 |
| FAK Y576/577 * | PI3K/PLCg | CellSig | 3281 | 1:1000 |

| | | | | |
|---|---|---|---|---|
| FGF Rec Y653/654 | PLCg | CellSig | 3471 | 1:1000 |
| FKHR/FKHRL1 T24/32 * | PI3 kinase/AKT | CellSig | 9464 | 1:1000 |
| FKHRL1 S253 * | PI3 kinase/AKT | Upstate | 06-953 | 1:500 |
| FOX01 S256 * | PI3 kinase/AKT | CellSig | 9461 | 1:100 |
| FOXM1 T600 * | PI3 kinase/AKT | CellSig | 14655 | 1:100 |
| GSK-3a/B S21/9 * | PI3 kinase/AKT | CellSig | 9331 | 1:1000 |
| HDAC4 * | Notch | CellSig | 2072 | 1:1000 |
| HDAC6 * | Notch | SantaCruz | sc-11420 | 1:2000 |
| Histone H3 * | p38 MAPK/Ras/PI3 kinase | Abcam | ab1791 | 1:500 |
| Histone H3 S10 * | p38 MAPK/Ras/PI3 kinase | Upstate | 06-570 | 1:1000 |
| Histone H3 S28 * | p38 MAPK/Ras/PI3 kinase | Upstate | 07-145 | 1:1000 |
| IGF-1 Rec/Insulin Rec Y1131/1146 * | PI3 kinase/AKT/Ras/MAPK | CellSig | 3021 | 1:1000 |
| IGF-1 Rec/Insulin Rec Y1135/36/Y1150/51 * | PI3 kinase/AKT/Ras/MAPK | CellSig | 3024 | 1:1000 |
| IL-11 * | Stat/Ras/Erk/PI3 kinase | SantaCruz | sc-7924 | 1:200 |
| Insulin Rec B* | PI3 kinase/MEK/ERK/p38 MAPK | CellSig | 3025 | 1:1000 |
| IRS-1 S612 | JNK/NFkB/PKC/mTOR | CellSig | 2386 | 1:1000 |
| Jak2 Y1007 * | Stat/PI3 kinase/AKT/MEK/ERK | CellSig | 3771 | 1:1000 |
| Ki67 (MIB-1) | Any growth regulator | DAKO | M7240 | 1:1000 |
| LKB1 S334 * | AMPK/PTEN/AKT | CellSig | 3055 | 1:1000 |
| LKB1 S428 * | AMPK/PTEN/AKT | CellSig | 3051 | 1:1000 |
| M-CSF Rec Y723 | PI3 kinase/PLCg | CellSig | 3155 | 1:100 |
| MARCKS S152/156 * | PKC | CellSig | 2741 | 1:1000 |
| MDM2 S166 * | p53/AKT | CellSig | 3521 | 1:1000 |
| MEK 1/2 S217/221 | MEK/ERK | CellSig | 9121 | 1:1000 |
| Met Y1234/1235 | PI3 kinase | CellSig | 3126 | 1:200 |
| MSK1 S360 | ERK/p38 MAPK | CellSig | 9594 | 1:1000 |
| NPM T199 * | Cell cycle | CellSig | 3541 | 1:1000 |
| p27 Kip1 | Cell cycle | BD | 610241 | 1:5000 |
| p27 T187 | Cell cycle | Zymed | 71-7700 | 1:50 |
| p38 MAPK T180/Y182 * | p38 MAPK | CellSig | 9211 | 1:1000 |
| p70 S6 Kin S371 | PI3 kinase | CellSig | 9208 | 1:1000 |
| p70 S6 Kin T389 | PI3 kinase | CellSig | 9205 | 1:1000 |
| p70 S6 Kin T412 | PI3 kinase | Upstate | 07-018 | 1:1000 |
| p70 S6 Kinase | PI3 kinase | CellSig | 9202 | 1:1000 |
| p90RSK S380 * | PI3 kinase/p38 MAPK/ERK | CellSig | 9341 | 1:1000 |
| p90RSK T359/S363 * | PI3 kinase/p38 MAPK/ERK | CellSig | 9344 | 1:200 |
| PAK 1/2 S199/204/S192/197 * | MAPK/AKT/MEK/ERK | CellSig | 2605 | 1:1000 |
| Paxillin Y118 | FAK | CellSig | 2541 | 1:1000 |
| PDGF Rec B | MAPK/PI3 kinase | Upstate | 06-498 | 1:500 |
| PDGF Rec B Y751 | MAPK/PI3 kinase | CellSig | 3161 | 1:1000 |
| PDK1 S241 * | AKT/PKC | CellSig | 3061 | 1:1000 |
| PIAS1 * | STAT/p53 | CellSig | 3550 | 1:1000 |

| | | | | |
|---|---|---|---|---|
| PKA C T197 * | AKT/PKC | CellSig | 4781 | 1:1000 |
| PKC a S657 | PKC | Upstate | 06-822 | 1:500 |
| PKC a/BII T638/641 | PKC | CellSig | 9375 | 1:1000 |
| PKC delta T505 | PKC | CellSig | 9374 | 1:1000 |
| PKC theta T538 | PKC | CellSig | 9377 | 1:1000 |
| PKC zeta/lambda T410/403 | PKC | CellSig | 9378 | 1:1000 |
| PLK1 T210 | Cell cycle regulators | BD | 558400 | 1:10,000 |
| PP2A a Subunit * | AKT/BAD/MEK/ERK/JAK/p53 | CellSig | 2039 | 1:1000 |
| PRAS40 T246 * | PI3 kinase/AKT/Autophagy | BioSource | 44-1100 | 1:1000 |
| Promyelocytic Leukemia Protein * | p53 | Bethyl | A301-167A | 1:000 |
| PTEN * | AKT/PKB/p53 | CellSig | 9552 | 1:1000 |
| PTEN S380 * | AKT/PKB/p53 | CellSig | 9551 | 1:1000 |
| Pyk2 Y402 | MAPK/PKC/FAK | CellSig | 3291 | 1:1000 |
| Raf S259 | MEK/MAPK/AKT | CellSig | 9421 | 1:500 |
| Ras-GRF1 * | PKA/MEK/ERK | CellSig | 3322 | 1:1000 |
| Ras-GRF1 S916 * | PKA/MEK/ERK | CellSig | 3321 | 1:1000 |
| Rb S780 | Cell cycle regulators | CellSig | 3590 | 1:1000 |
| Ret Y905 | STAT/ERK | CellSig | 3221 | 1:1000 |
| Ron Y1353 | PI3 kinase/AKT | Epitomics | 5176-1 | 1:1000 |
| RSK3 T356/S360 * | MAPK/ERK/PI3 kinase/NFkB | CellSig | 9348 | 1:1000 |
| S6 Ribosomal Protein S235/236 | PI3 kinase/AKT | CellSig | 4856 | 1:1000 |
| S6 Ribosomal Protein S240/244 | PI3 kinase/AKT | CellSig | 2215 | 1:2000 |
| SAPK/JNK T183/Y185 * | MAPK/MEK/Rho | CellSig | 9251 | 1:1000 |
| SEK1/MKK4 S80 | p38 MAPK/ERK/AKT | CellSig | 9155 | 1:500 |
| SGK1 S78 * | PI3 kinase/p53/PKC | CellSig | 5599 | 1:500 |
| Shc Y317 * | MAPK/ERK | Upstate | 07-206 | 1:1000 |
| Smad2 (S465/467)* | TGFB | CellSig | 3101 | 1:1000 |
| Smad2 (S245/250/255) * | TGFB | CellSig | 3104 | 1:1000 |
| Src Family Y416 | MEK/STAT/Rho/PI3 kinase/AKT | CellSig | 2101 | 1:1000 |
| Src Y527 | MEK/STAT/Rho/PI3 kinase/AKT | CellSig | 2105 | 1:1000 |
| Stat5 Y694 * | Stat | CellSig | 9351 | 1:1000 |
| Survivin * | p53/Wnt | CellSig | 2808 | 1:1000 |
| Syk Y525/526 * | PLCg/PI3 kinase/AKT/MAPK/NFkB | CellSig | 2711 | 1:1000 |
| TGF-B * | MEK/ERK/PI3 kinase/ AKT/Rho/MAPK | CellSig | 3709 | 1:1000 |
| Tubulin a * | | Sigma | T 6074 | 1:4000 |
| Tyk2 Y1054/1055 * | MAPK/Stat | CellSig | 9321 | 1:1000 |
| VASP S157 | PKA/PKG | CellSig | 3111 | 1:1000 |
| YAP S127 * | Akt/PI3 kinase/Hippo | CellSig | 13008 | 1:100 |
| Zap-70/Syk Y319/352 * | PLCg/PI3 kinase/AKT/MAPK/NFkB | CellSig | 2701 | 1:500 |

| **Protein Name** | **Pathways** **Transcription/Translation Regulators** | **Company** | **Catalog #** | **Dilutions** |
|---|---|---|---|---|

| | | | | |
|---|---|---|---|---|
| 4E-BP1 S65 | Down stream PI3 kinase/ Akt and FRAP/mTOR | CellSig | 9451 | 1:1000 |
| 4E-BP1 T70 | Down stream PI3 kinase/ Akt and FRAP/mTOR | CellSig | 9455 | 1:1000 |
| Acetyl-Histone H3 Lys9/14 | Gene regulation/p38 MAPK | CellSig | 9677 | 1:2000 |
| Acetyl-Histone H4 Lys8 | Gene regulation/ p38 MAPK | CellSig | 2594 | 1:200 |
| Di-Methyl Histone H3 Lys9 | PRMT signalling/p38 MAPK | CellSig | 9753 | 1:500 |
| eIF4E S209 | PI3 kinase/AKT/mTOR/Ras/MAPK | CellSig | 9741 | 1:1000 |
| eIF4G S1108 | PI3 kinase/AKT/mTOR/Ras/MAPK | CellSig | 2441 | 1:1000 |
| Elk-1 S383 | p38 MAPK | CellSig | 9181 | 1:1000 |
| HDAC4 * | Notch | CellSig | 2072 | 1:1000 |
| HDAC6 * | Notch | SantaCruz | sc-11420 | 1:2000 |
| Histone H3 * | p38 MAPK/Ras/PI3 kinase | Abcam | ab1791 | 1:500 |
| Histone H3 S10 * | p38 MAPK/Ras/PI3 kinase | Upstate | 06-570 | 1:1000 |
| Histone H3 S28 * | p38 MAPK/Ras/PI3 kinase | Upstate | 07-145 | 1:1000 |
| NPM T199 * | Cell cycle | CellSig | 3541 | 1:1000 |
| Pan-Methyl Histone H3 Lys9 | PRMT signalling/p38 MAPK | CellSig | | 1:100 |
| **Protein Name** | **Pathways** | **Company** | **Catalog #** | **Dilutions** |
| | **Apoptosis** | | | |
| BAD | AKT | CellSig | 9292 | 1:1000 |
| BAD S112 | AKT | CellSig | 9291 | 1:1000 |
| BAD S136 | AKT | CellSig | 9295 | 1:500 |
| PKA C T197 * | AKT/TNF | CellSig | 4781 | 1:1000 |
| Bak | AKT? | CellSig | 3814 | 1:1000 |
| Bax | p53 | CellSig | 2772 | 1:1000 |
| Bcl-2 | p53? | CellSig | 2872 | 1:1000 |
| Bcl-2 S70 | p53? | CellSig | 2827 | 1:1000 |
| Beclin 1 * | PI3 kinase | Sigma | PRS3613 | 1:500 |
| BIM | AKT/p38 MAPK/MEK;ERK | CellSig | 2933 | 1:1000 |
| c-Abl T735 * | p53 | CellSig | 2864 | 1:50 |
| c-Met * | Upstream RAS/PI3K/ STAT/Mnt/Notch | Abcam | ab51067 | 1:200 |
| c-Myc * | p38 MAPK/ERK | CellSig | 9402 | 1:100 |
| c-Src * | AKT/PI3 kinase/p38 MAPK/ MEK;ERK | SantaCruz | sc-18 | 1:200 |
| Cleaved Caspase-7 D198 | TNF | CellSig | 9491 | 1:1000 |
| Cleaved Caspase-9 D330 | TNF | CellSig | 9501 | 1:1000 |
| Cleaved PARP D214 | TNF | CellSig | 9541 | 1:100 |
| ErbB2 Y1248 * | PI3 kinase/AKT/PLC/ PKC/MAPK/STAT | CellSig | 2247 | 1:1000 |
| ErbB2 Y877 * | PI3 kinase/AKT/PLC/ PKC/MAPK/STAT | Imgenex | IMG-90189 | 1:500 |
| ErbB3 * | PI3 kinase/AKT | CellSig | 4754 | 1:1000 |
| ErbB3 Y1289 * | PI3 kinase/AKT | CellSig | 4791 | 1:1000 |
| ERK 1/2 T202/Y204 * | MAPK | CellSig | 9101 | 1:1000 |

| | | | | |
|---|---|---|---|---|
| FADD S194 | TNF | CellSig | 2781 | 1:1000 |
| FAK Y576/577 * | PI3K/PLCg | CellSig | 3281 | 1:1000 |
| FKHR/FKHRL1 T24/32 * | PI3 kinase/AKT | CellSig | 9464 | 1:1000 |
| FKHRL1 S253 * | PI3 kinase/AKT | Upstate | 06-953 | 1:500 |
| FOX01 S256 * | PI3 kinase/AKT | CellSig | 9461 | 1:100 |
| FOXM1 T600 * | PI3 kinase/AKT | CellSig | 14655 | 1:100 |
| GSK-3a/B S21/9 * | PI3 kinase/AKT | CellSig | 9331 | 1:1000 |
| HDAC4 * | Notch | CellSig | 2072 | 1:1000 |
| HDAC6 * | Notch | SantaCruz | sc-11420 | 1:2000 |
| HSP27 Protein 1 | p38 MAPK/ERK/NFkB | Lifespan Bio | LS-C34438 | 1:500 |
| IGF-1 Rec/Insulin Rec Y1131/1146 * | PI3 kinase/AKT/Ras/MAPK | CellSig | 3021 | 1:1000 |
| IGF-1 Rec/Insulin Rec Y1135/36/Y1150/51 * | PI3 kinase/AKT/Ras/MAPK | CellSig | 3024 | 1:1000 |
| Jak2 Y1007 * | Stat/PI3 kinase/AKT/MEK/ERK | CellSig | 3771 | 1:1000 |
| LKB1 S334 * | AMPK/PTEN/AKT | CellSig | 3055 | 1:1000 |
| LKB1 S428 * | AMPK/PTEN/AKT | CellSig | 3051 | 1:1000 |
| MDM2 S166 * | p53/AKT | CellSig | 3521 | 1:1000 |
| p90RSK S380 * | PI3 kinase/p38 MAPK/ERK | CellSig | 9341 | 1:1000 |
| p90RSK T359/S363 * | PI3 kinase/p38 MAPK/ERK | CellSig | 9344 | 1:200 |
| PAK 1/2 S199/204/S192/197 * | MAPK/AKT/MEK/ERK | CellSig | 2605 | 1:1000 |
| PDK1 S241 * | AKT/PKC | CellSig | 3061 | 1:1000 |
| PIAS1 * | STAT/p53 | CellSig | 3550 | 1:1000 |
| PKA C T197 * | AKT/PKC | CellSig | 4781 | 1:1000 |
| PP2A a Subunit * | AKT/BAD/MEK/ERK/JAK/p53 | CellSig | 2039 | 1:1000 |
| PRAS40 T246 * | PI3 kinase/AKT/Autophagy | BioSource | 44-1100 | 1:1000 |
| Promyelocytic Leukemia Protein * | p53 | Bethyl | A301-167A | 1:000 |
| PTEN * | AKT/PKB/p53 | CellSig | 9552 | 1:1000 |
| PTEN S380 * | AKT/PKB/p53 | CellSig | 9551 | 1:1000 |
| Ras-GRF1 * | PKA/MEK/ERK | CellSig | 3322 | 1:1000 |
| Ras-GRF1 S916 * | PKA/MEK/ERK | CellSig | 3321 | 1:1000 |
| SAPK/JNK T183/Y185 * | MAPK/MEK/Rho | CellSig | 9251 | 1:1000 |
| SGK1 S78 * | PI3 kinase/p53/PKC | CellSig | 5599 | 1:500 |
| Shc Y317 * | MAPK/ERK | Upstate | 07-206 | 1:1000 |
| Smad2 (S465/467) * | TGFB | CellSig | 3101 | 1:1000 |
| Smad2 (S245/250/255) * | TGFB | CellSig | 3104 | 1:1000 |
| Stat5 Y694 * | Stat | CellSig | 9351 | 1:1000 |
| Stat6 Y641 * | Stat/NFkB | CellSig | 9361 | 1:1000 |
| Survivin * | p53/Wnt | CellSig | 2808 | 1:1000 |
| Syk Y525/526 * | PLCg/PI3 kinase/AKT/MAPK/NFkB | CellSig | 2711 | 1:1000 |
| TGF-B * | MEK/ERK/PI3 kinase/ AKT/ Rho/MAPK | CellSig | 3709 | 1:1000 |
| TNF a * | MEK/MAPK/NFkB | Abcam | ab9635 | 1:500 |
| TNF-R1 * | MEK/MAPK/NFkB | CellSig | 3736 | 1:500 |
| Tyk2 Y1054/1055 * | MAPK/Stat | CellSig | 9321 | 1:1000 |

| | | | | |
|---|---|---|---|---|
| YAP S127 * | Akt/PI3 kinase/Hippo | CellSig | 13008 | 1:100 |
| XIAP | TNF/NFkB | CellSig | 2042 | 1:1000 |
| Zap-70/Syk Y319/352 * | PLCg/PI3 kinase/AKT/MAPK/NFkB | CellSig | 2701 | 1:500 |
| **Protein Name** | **Pathways** | **Company** | **Catalog #** | **Dilutions** |
| | **Immune Response** | | | |
| CD3 epsilon | T-cell activation | Abcam | Ab5690 | 1:1000 |
| CrkL Y207 * | Ras/Jun/PI3 kinase | CellSig | 3181 | 1:1000 |
| Cox-2 | p38 MAPK/MEK;ERK | BD | 610203 | 1:200 |
| eNOS S113 * | AKT/PKA/AMPK | CellSig | 9575 | 1:1000 |
| eNOS/NOSIII S116 * | AKT/PKA/AMPK | Upstate | 07-357 | 1:1000 |
| Ephrin A3/A4/A5 Y799/799/833 * | Ephrin/Rac | Abcam | 124881 | 1:100 |
| IkBa S32/36 | NFkB | CellSig | 9246 | 1:2000 |
| IL-8 | Stat/Ras/Erk/PI3 kinase/AKT | Abcam | ab7747 | 1:400 |
| IL-11 * | Stat/Ras/Erk/PI3 kinase | SantaCruz | sc-7924 | 1:200 |
| Insulin Rec B * | PI3 kinase/MEK/ERK/p38 MAPK | CellSig | 3025 | 1:1000 |
| Jak1 Y1022/1023 | Stat/IFN | CellSig | 3331 | 1:1000 |
| Lck Y505 | PI3 kianse/PLC/MEK/ERK/NFkB | Biosource | 44-850 | 1:1000 |
| Lysine 48-Linkage Specific Polyubiquitin * | NFkB/p53/autophagy | CellSig | 4289 | 1:1000 |
| MARCKS S152/156 * | PKC | CellSig | 2741 | 1:1000 |
| NF-kB S536 | NFkB/p38 MAPK | CellSig | 3031 | 1:1000 |
| p38 MAPK T180/Y182 * | p38 MAPK | CellSig | 9211 | 1:1000 |
| p62/SQSTM1 * | PKC/NFkB/Autophagy | MBL | M162-3B | 1:1000 |
| PAK 1/2 S199/204/S192/197 * | MAPK/AKT/MEK/ERK | CellSig | 2605 | 1:1000 |
| PD-L1 | PKC NFkB | CellSig | 13684 | 1:500 |
| PIAS1 * | STAT/p53 | CellSig | 3550 | 1:1000 |
| Promyelocytic Leukemia Protein * | p53 | Bethyl | A301-167A | 1:000 |
| RSK3 T356/S360 * | MAPK/ERK/PI3 kinase/NFkB | CellSig | 9348 | 1:1000 |
| SAPK/JNK T183/Y185 * | MAPK/MEK/Rho | CellSig | 9251 | 1:1000 |
| SGK1 S78 * | PI3 kinase/p53/PKC | CellSig | 5599 | 1:500 |
| Shc Y317 * | MAPK/ERK | Upstate | 07-206 | 1:1000 |
| Stat1 Y701 | p38 MAPK/Stat | CellSig | 9171 | 1:1000 |
| Stat2 Y690 | MAPK/Stat | CellSig | 4441 | 1:1000 |
| Stat4 Y693 | Stat | CellSig | 5267 | 1:1000 |
| Stat5 Y694 * | Stat | CellSig | 9351 | 1:1000 |
| Stat6 Y641 * | Stat/NFkB | CellSig | 9361 | 1:1000 |
| Syk Y525/526 * | PLCg/PI3 kinase/AKT/MAPK/NFkB | CellSig | 2711 | 1:1000 |
| TNF a * | MEK/MAPK/NFkB | Abcam | ab9635 | 1:500 |
| TNF-R1 * | MEK/MAPK/NFkB | CellSig | 3736 | 1:500 |
| Tyk2 Y1054/1055 * | MAPK/Stat | CellSig | 9321 | 1:1000 |
| Zap-70/Syk Y319/352 * | PLCg/PI3 kinase/AKT/MAPK/NFkB | CellSig | 2701 | 1:500 |
| **Protein Name** | **Pathways** | **Company** | **Catalog #** | **Dilutions** |
| | **Autophagy** | | | |
| ATG5 | mTOR/AMPK | CellSig | 2630 | 1:500 |

| | | | | |
|---|---|---|---|---|
| Beclin 1 * | PI3 kinase | Sigma | PRS3613 | 1:500 |
| LAMP-2 | Autophagy | SantaCruz | sc-18822 | 1:200 |
| LC3B | Autophagy | CellSig | 2775 | 1:500 |
| p62/SQSTM1 * | PKC/NFkB/Autophagy | MBL | M162-3B | 1:1000 |
| PRAS40 T246 * | PI3 kinase/AKT/Autophagy | BioSource | 44-1100 | 1:1000 |

| **Protein Name** | **Pathways** | **Company** | **Catalog #** | **Dilutions** |
|---|---|---|---|---|
| | **Cell Structure/Metabolism** | | | |
| Acetyl-CoA Carboxylase S79 | Fatty acid | CellSig | 3661 | 1:500 |
| Actin, Beta | Cytoskeleton | CellSig | 4967 | 1:1000 |
| ATP-Citrate Lyase S454 | AKT/PKA | CellSig | 4331 | 1:1000 |
| Catenin B S33/37/T41 | Wnt | CellSig | 9561 | 1:1000 |
| Cofilin S3 | Rac | CellSig | 3313 | 1:1000 |
| CrkL Y207 * | Ras/Jun/PI3 kinase | CellSig | 3181 | 1:1000 |
| Estrogen Rec a S118 | MAPK/PI3 kinase/AKT | CellSig | 2515 | 1:500 |
| Ezrin/Radixin/Moesin T567/564/558 * | Rho | CellSig | 3141 | 1:1000 |
| LIMK 1/2 T508/505 | Rho | CellSig | 3841 | 1:1000 |
| Lysine 48-Linkage Specific Polyubiquitin * | NFkB/p53/autophagy | CellSig | 4289 | 1:1000 |
| MMP-11 | MEK/ERK/PKA | Abcam | ab52904 | 1:1000 |
| MMP-9 | MEK/ERK/PKA | CellSig | 3852 | 1:500 |
| Tubulin a* | | Sigma | T 6074 | 1:4000 |
